# Role of aIF5B in archaeal translation initiation

**DOI:** 10.1101/2022.05.01.490067

**Authors:** Ramy Kazan, Gabrielle Bourgeois, Christine Lazennec-Schurdevin, Eric Larquet, Yves Mechulam, Pierre-Damien Coureux, Emmanuelle Schmitt

**Affiliations:** Laboratoire de Biologie Structurale de la Cellule, BIOC, Ecole polytechnique, CNRS, Institut Polytechnique de Paris, 91128 Palaiseau cedex, France; Laboratoire de Physique de la Matière Condensée, PMC, Ecole polytechnique, CNRS, Institut Polytechnique de Paris, 91128 Palaiseau cedex, France

**Author notes:** Corresponding authors: Emmanuelle Schmitt, Pierre-Damien Coureux, Yves Mechulam,.

**Keywords:** ribosome, initiator tRNA, eIF5B, cryo-EM

## Abstract

In eukaryotes and in archaea late steps of translation initiation involve the two initiation factors e/aIF5B and e/aIF1A. In eukaryotes, the role of eIF5B in ribosomal subunit joining is established and structural data showing eIF5B bound to the full ribosome were obtained. To achieve its function, eIF5B collaborates with eIF1A. However, structural data illustrating how these two factors interact on the small ribosomal subunit have long been awaited. The role of the archaeal counterparts, aIF5B and aIF1A, remains to be extensively addressed. Here, we study the late steps of *Pyrococcus abyssi* translation initiation. Using *in vitro* reconstituted initiation complexes and light scattering, we show that aIF5B bound to GTP accelerates subunit joining without the need for GTP hydrolysis. We report the crystallographic structures of aIF5B bound to GDP and GTP and analyze domain movements associated to these two nucleotide states. Finally, we present the cryo-EM structure of an initiation complex containing 30S bound to mRNA, Met-tRNA_i_^Met^, aIF5B and aIF1A at 2.7 Å resolution. Structural data shows how archaeal 5B and 1A factors cooperate to induce a conformation of the initiator tRNA favorable to subunit joining. Archaeal and eukaryotic features of late steps of translation initiation are discussed.

## INTRODUCTION

Studies of the central dogma processes, such as protein biosynthesis, allow better understanding of evolution. Thus, pioneering analysis of small ribosomal subunit RNA led to define the three-domain tree of life with archaea close to eucaryotes (1,2). Recent phylogenomic studies based on new methods and on the discovery of new archaeal species proposed a two-domain tree of life, bacteria and archaea, with eukaryotes that could have emerged from within an archaeal branch (3-6). In these models, archaea appear as a central piece to understand evolution of life and efforts to better understand their diversity and molecular mechanisms will bring important insights.

In archaea, mRNAs are not maturated after their transcription and present Shine-Dalgarno sequences or very short 5’ untranslated regions, as in bacteria. However, other features of the translation apparatus bring archaea closer to eukaryotes. In particular, the archaeal ribosome and translation factors are of the eukaryotic type (7-17).

Archaeal translation initiation (TI) is divided into two stages, as in eukaryotes (10,18). The early steps of TI are devoted to start codon selection and were recently illustrated in the case of *Pyrococcus abyssi* by cryo-EM structures (19-21). Start codon selection is achieved within a macromolecular complex involving the small ribosomal subunit (SSU) bound to the mRNA, aIF1, aIF1A and the ternary complex aIF2:GTP:Met-tRNA_i_^Met^. aIF2 is a heterotrimeric protein (α, β, γ subunits) that binds GTP and the methionylated initiator tRNA (22,23). aIF1A occupies the A site and aIF1 is located in front of the P site. The current model suggests that the interaction of the start codon with the initiator tRNA anticodon triggers the departure of aIF1 and then that of aIF2:GDP. aIF2 and aIF1 have an important role for the accuracy of start codon selection (20,24-26). Because aIF1, aIF1A and aIF2 are homologous to their eukaryotic counterparts, selection of the start codon in eukaryotes and in archaea is carried out within a common structural core (21).

In eukaryotes and in archaea, late steps of translation initiation occur after start codon selection and e/aIF2 departure. These steps involve the two initiation factors e/aIF1A and e/aIF5B (9,27,28). Importantly, e/aIF5B and e/aIF1A are orthologues of the bacterial proteins IF2 and IF1, respectively. Late steps of translation initiation have therefore a universal character (29). In eukaryotes, these two factors ensure the final checkpoint for the presence of the initiator tRNA on the SSU and facilitate the assembly with the large ribosomal subunit (LSU) (27,30-32). Subsequent GTP hydrolysis-dependent departure of eIF5B enables protein synthesis to begin (30,33-35). e/aIF5B is composed of four domains (36-40). Domains I (GTP binding domain), II and III are packed together and linked by a long α–helix (helix h12) to domain IV responsible for the binding of the methionylated-CCA end of Met-tRNA_i_^Met^ (41). Eukaryotic eIF5Bs contain an additional N-domain with little sequence conservation that was shown to be dispensable in yeast (33). Several cryo-EM structures of 80S:eIF5B complexes were determined (42-46). These structures show in particular that eIF5B-domain IV is bound to the Met-CCA end of the initiator tRNA and that domain I interacts with the sarcin-ricin region of the LSU. As for other translation GTPases, the nucleotide cycle is controlled by two switch regions that are either in ON state when a/aIF5B is bound to GTP or in OFF state when the factor is bound to GDP (37). In eukaryotes, the integrity of the h12 helix and the multidomain nature of the factor were shown important for its function (37,43,47). Finally, eIF5B was shown to directly interact with the eukaryote-specific C-terminal tail of eIF1A (40,48,49). This interaction is required for optimal subunit joining (35,50). However, until very recently, this interaction had not been observed because structural data of TI complexes on the small ribosomal subunit were still missing.

In archaea, aIF1A does not possess the eukaryotic-specific C-terminal extension but aIF5B possesses a supplementary C-terminal helix (39). Therefore, the possibility that the two proteins interact on the small ribosomal subunit remains to be explored. Moreover, even if previous work gave insights into the function of aIF5B (28,38,39,41,51,52), the role of aIF5B in the late steps of TI remains to be extensively studied.

In the present work, we study the *P. abyssi* (Pa) late steps of translation initiation. Using *in vitro* reconstitutions of translation initiation complexes and light scattering, we show that the transition from a 30S preinitiation complex containing aIF2 to the final 70S initiation complex is accelerated by aIF5B bound to GTP. GTP hydrolysis is not required for the joining step. We report the crystallographic structures of *P. abyssi* aIF5B bound to GDP and GTP and analyze domain movements associated to these two nucleotide states. Finally, we present the cryo-EM structure of an initiation complex containing 30S bound to mRNA, Met-tRNA_i_^Met^, aIF5B and aIF1A at 2.7 Å resolution. The structure shows how the archaeal aIF5B and aIF1A factors cooperate to induce a conformation of the initiator tRNA favorable to subunit joining.

While writing our results, two manuscripts describing mammalian complexes containing the 40S subunit, eIF5B and eIF1A were deposited in BioRxiv (53,54). Our data will also be discussed at the light of these recent results.

## MATERIALS AND METHODS

### Production and purification of aIF5B and variants

The *P. abyssi* gene coding for mature (intein free) aIF5B was subcloned into pET15blpa from pET3a5Bpa (41). The resulting plasmid pET15blpa-Pa-aIF5B encodes an N-terminally His_6_-tagged version of the mature aIF5B protein. Rosetta pLacI-RARE cells (Novagen) carrying pET15blpa-Pa-aIF5B were grown at 37°C in 1 L of 2xTY medium containing ampicillin (50 µg/mL) and chloramphenicol (34 µg/mL). Expression was induced by adding IPTG to a final concentration of 1 mM when OD_600nm_ reached 2. The culture was then continued for 5 h at 18°C. Cells were disrupted by sonication in 40 mL of buffer A (10 mM MOPS pH 6.7, 200 mM NaCl, 3 mM 2-mercaptoethanol, 0.1 mM PMSF, 0.1 mM benzamidine). After centrifugation, the supernatant was heated for 10 min at 75°C. The precipitated material was removed by centrifugation and the soluble fraction was loaded onto a column (5 mL) containing Talon affinity resin (Clontech) equilibrated in buffer A. After washing the column with buffer A supplemented with 10 mM imidazole, the tagged protein was eluted with buffer A supplemented with 125 mM imidazole. The pooled fractions were then loaded onto an S-sepharose column (5 mL; Cytiva) and eluted with an NaCl gradient. The homogeneous protein was concentrated to 3 mg/mL by using a Centricon-30 concentrator. Nearly 8 mg of protein per liter of culture was obtained.

Deletion of the C-terminal domain and of part of h12 helix (from residue 459 to residue 598) was obtained via mutagenesis of plasmid pET5blpa-Pa-aIF5B according to the QuikChange™ site-directed mutagenesis method (Stratagene). This deletion was designed after inspection of known structures (37) and of the structure of full-length aIF5B determined in the present study. Overexpression and purification of the variant were carried out as for the full-length protein. The shortened protein is hereafter named aIF5B-ΔC. H82A and Y440A variants were produced using the QuikChange™ site-directed mutagenesis method (Stratagene) and purified as wild-type aIF5B.

### Other materials

Pa-30S subunits were purified as described (19,20). Pa-50S subunits were purified using the same strategy. After 10-30 % sucrose gradient fractionation, fractions containing 30S and those containing 50S subunits were pooled separately and concentrated. To improve the purity of the 50S subunits, a second 10-40 % sucrose gradient fractionation was performed (Beckmann SW32.1 rotor, 220,000 g, 20 h). 50S containing fractions were recovered and their quality was analyzed by SDS and native gel electrophoresis. *E. coli* tRNA_f_^Met^ A_1_ -U_72,_ hereafter named tRNA_i_^Met^ was produced in *E. coli* from a cloned gene, purified and aminoacylated as described (55). This tRNA was shown previously to be an effective mimic of the *P. abyssi* initiator tRNA (19,20,56-58). A synthetic 26 nucleotide long mRNA corresponding to the natural start region of the mRNA encoding elongation factor aEF1A from *P. abyssi*, which contains a strong Shine-Dalgarno sequence (A_(- 17)_UUU**GGAGGUGAU**UUAA**A**_(+1)_**UG**CCAAAG_(+9)_, Dharmacon) was used as in (19). The gene encoding aIF1A from *P. abyssi* was amplified from genomic DNA and cloned into pET15blpa to produce an N-terminally his-tagged version of the factor. aIF1A was purified as described (24) except that a standard affinity chromatography step on Talon resin (Clontech) was added to the original protocol. *P. abyssi* aIF2 and aIF1 were purified as described (20).

### GTP hydrolysis assay

GTP hydrolysis was monitored in reactions containing 6.6 nM γ_32_[P]-GTP (Hartmann Analytic, Germany) in 10mM MOPS-NaOH pH 6.7, 100 mM NH_4_Cl, 10.5 mM Mg acetate, 2 mM 2-mercaptoethanol, 10 mM potassium phosphate (pH 7.0), and the following components when indicated (200 nM 30S subunits, 200 nM aIF1A, 200 nM synthetic mRNA, 200 nM Met-tRNA_i_^Met^, Table 1). Prior to the assay, aIF5B and derivatives were dialyzed against 10 mM MOPS pH 6.7, 200 mM NaCl, 5mM 2-mercaptoethanol, supplemented with 2 mM MgCl_2_ at 65°C during 3 hours and ribosomal subunits were activated at least 30 minutes at 51°C. aIF5B or aIF5BΔC was added at an optimal concentration of 1 µM (0.5 µM for the Y440A variant). The 15 µL reactions were started by adding 50S subunits (200 nM final concentration) and incubated at 51°C in a thermal cycler apparatus to avoid evaporation. 1.5 µL aliquots were withdrawn at various times and quenched by adding 1 mL of a suspension containing 0.4 % norite, 0.35 % perchloric acid, 50 mM sodium acetate pH 5.0, 100 mM pyrophosphate. The remaining labelled γ^32^[P]-GTP was adsorbed on norite, recovered by filtration on Whatman filter paper disks, washed with water and counted on a Perkin-Elmer Tri-Carb 4910TR scintillation counter. The rates of GTP hydrolysis were determined from iterative non-linear least-square fits of the experimental points to single decreasing exponentials (Figure 1A and Table 1).

**Table 1:** GTP hydrolysis rate constants. Rates were measured at 51°C as described in the methods section. *With aIF5B and 50S subunits, slow hydrolysis (k=0.055 ± 0.002 min^-1^) was observed. However, this hydrolysis is likely due to contamination of 50S by 30S. Each experiment was repeated 4-5 times from which rate constants (k) and associated standard deviation were calculated.

| Components | Rate constant (min<sup>-1</sup>) |
| --- | --- |
| 1-aIF5B alone | <0.01 |
| 2- aIF5B + 30S | <0.01 |
| 3- aIF5B + 50S* | 0.055 ± 0.002 |
| 4-aIF5B:30S + 50S | 0.25 ± 0.02 |
| 5-aIF5B:30S:mRNA:Met-tRNA<sub>i</sub><sup>Met</sup> + 50S | 0.35 ± 0.04 |
| 6-aIF5B:30S:mRNA:Met-tRNA<sub>i</sub><sup>Met</sup>:aIF1A + 50S | 0.32 ± 0.02 |
| 7-aIF5B-H82A:30S:mRNA:Met-tRNA<sub>i</sub><sup>Met</sup>:aIF1A + 50S | <0.01 |
| 8-aIF5B-Y440A:30S:mRNA:Met-tRNA<sub>i</sub><sup>Met</sup>:aIF1A + 50S | 0.33 ± 0.06 |
| 9-aIF5B-ΔC:30S:mRNA:Met-tRNA<sub>i</sub><sup>Met</sup>:aIF1A + 50S | 0.30 ± 0.06 |
| 10-aIF5B-ΔC:30S + 50S | 0.22 ± 0.03 |

**Figure 1:**
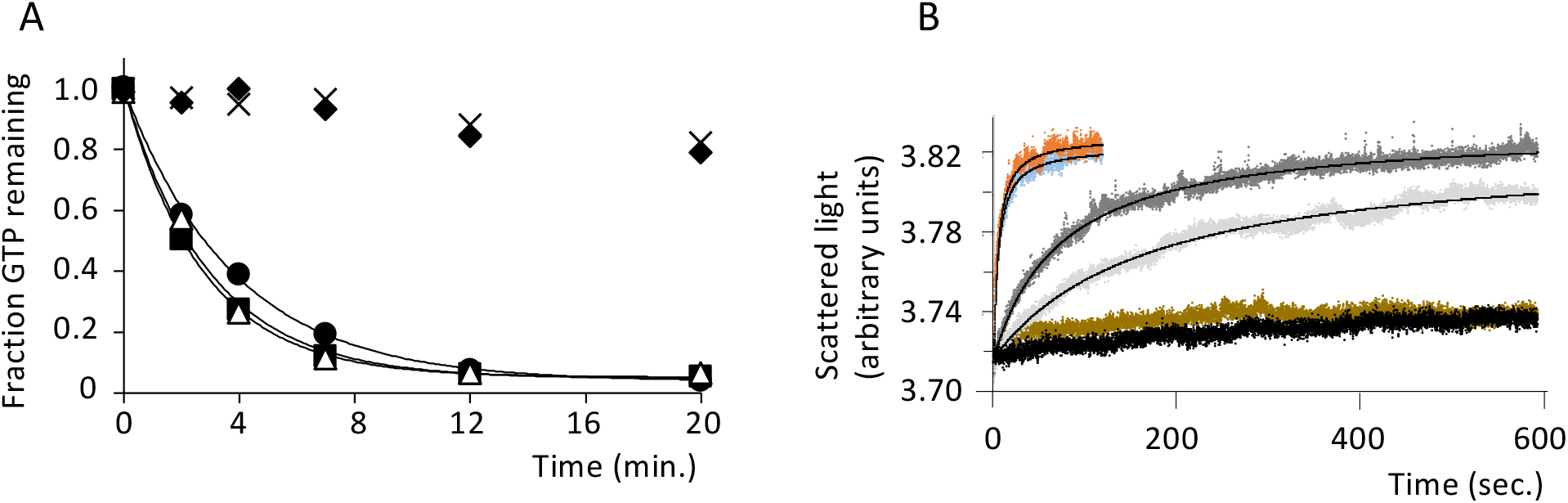
Pa-aIF5B is active in GTP hydrolysis and subunit joining. (**A**) Time course of [γ]^32^P-GTP hydrolysis by aIF5B. aIF5B (1 µM) was incubated at 51°C with [γ]^32^P-GTP (13 nM) in the presence of the indicated components. At the indicated times, the fraction of remaining GTP was measured (see Materials and Methods). Data points were fitted with single exponentials. Each experiment was repeated 4-5 times from which rate constants (k) and associated standard deviation were calculated. Representative experiments are shown. Added components were: 30S and 50S subunits (200 nM each, closed circles, k=0.25 ± 0.02 min^-1^), 30S, 50S, Met-tRNA_i_^Met^ and mRNA (200 nM each, closed squares, k=0.35 ± 0.04 min^-1^), 30S, 50S, Met-tRNA_i_^Met^, mRNA and aIF1A (200 nM each, open triangles, k=0.32 ± 0.02 min^-1^). The crosses represent the same experiment with 30S, 50S, Met-tRNA_i_^Met^, mRNA and aIF1A, but aIF5B was replaced by its H82A mutant (k < 0.01 min-1). A control experiment with aIF5B alone is represented by closed diamonds (k < 0.01 min-1). The same result (k < 0.01 min^-1^) was obtained in the presence of aIF5B and 30S subunits. With aIF5B and 50S subunits, slow hydrolysis (k=0.055 ±0.002 min^-1^) was observed that is probably due to some contamination of 50S subunits by 30S subunits. (**B**) Time course of ribosomal subunits association as measured by light scattering. Data points were fitted to a hyperbolic equation (61) from which rate constants were deduced. The figure shows representative experiments with the data points and the corresponding fitted curves. In the representation, data were scaled using the fitted value of the zero point in order to facilitate comparison. Black: 30S and 50S subunits assembled in the presence of GTP only; Blue: 30S subunits preincubated with aIF2, Met-tRNA_i_^Met^, mRNA, aIF1A and GTP, mixed with 50S subunits, aIF5B and GTP; Orange: same experiment but aIF5B was replaced by its H82A variant; Brown: 30S subunits preincubated with aIF2, Met-tRNA_i_^Met^, mRNA, aIF1A and GTP, mixed with 50S subunits and GTP; light grey: 30S subunits containing Met-tRNA_i_^Met^, mRNA, and GTP, mixed with 50S subunits and GTP; dark grey: 30S and 50S subunits assembled in the presence of aIF5B-H82A and GTP only. Rate constants deduced from the average of at least 3 measurements were 11.4 ± 3.0 min^- 1^ (blue); 11.4 ± 1.8 min^-1^ (orange); 0.32 ± 0.04 min^-1^ (light grey); 0.84 ± 0.06 min^-1^ (dark grey).

### Subunits association monitored by light scattering

The association of ribosomal subunits was monitored by light scattering after rapid mixing in an SX.18MV stopped-flow apparatus (Applied Photophysics, Leatherhead, UK), as previously described (32,59,60). The excitation wavelength was 435 nm (4.65 nm bandpass) and the scattered light was measured at an angle of 90° to the incident beam. Before mixing, the 30S and 50S subunits were incubated at least 30 min at 51°C. Two mixes were prepared as follows. The mix corresponding to Syringe 1 contained 30S ribosomal subunits (50 nM), GTP (250 µM) with or without, depending on the experiment, mRNA (50nM), Met-tRNA_i_^Met^ (50nM), aIF2-GTP (50nM), aIF1 (50nM) and aIF1A (50nM) (Table 2). Mix corresponding to Syringe 2 contained 50S ribosomal subunits (50 nM), GTP (250 µM) with or without, depending on the experiment, aIF5B or its variants (1 µM). In Table 2, lane 8, GDPNP (250 µM) was used instead of GTP in syringe 2. Association was measured at 51°C after at least 30 minutes of incubation to complete hydrolysis of GTP bound to aIF2 in Syringe 1 (see Results). The observed association rate constants were estimated by non-linear least-square fitting of the scattered intensity (I) data points to a hyperbolic equation (I=I_final_-D(1/1+k_obs_*t)) as described (61). The Origin (OriginLab) software was used for fitting (Figure 1B).

**Table 2:** Observed rate constants for ribosome assembly. A volume of components contained in syringe 1 (∼ 60 µL) is mixed to an equal volume of components contained in syringe 2 using a stopped-flow apparatus (Materials and Methods). Time course of ribosomal subunits association is followed by measuring light scattering. Data points were fitted to a hyperbolic equation (61) from which rate constants were deduced. n.m.: not measurable. Rate constants are deduced from the average of at least 3 measurements.

| Syringe 1 | Syringe 2 | Observed rate constant (min<sup>-1</sup>) |
| --- | --- | --- |
| 1-30S + GTP | 50S | n.m |
| 2-30S + GTP | 50S + aIF5B-H82A + GTP | 0.84 ± 0.06 |
| 3-30S:Met-tRNA<sub>i</sub><sup>Met</sup>:mRNA + GTP | 50S + GTP | 0.32 ± 0.04 |
| 4-30S:Met-tRNA<sub>i</sub><sup>Met</sup>:mRNA:aIF2:aIF1:aIF1A + GTP | 50S + GTP | n.m |
| 5-30S:Met-tRNA<sub>i</sub><sup>Met</sup>:mRNA:aIF2:aIF1:aIF1A + GTP | 50S + aIF5B + GTP | 11.4 ± 3.0 |
| 6-30S:Met-tRNA<sub>i</sub><sup>Met</sup>:mRNA:aIF2:aIF1:aIF1A + GTP | 50S + aIF5B-H82A + GTP | 11.4 ± 1.8 |
| 7-30S:Met-tRNA<sub>i</sub><sup>Met</sup>:mRNA:aIF2:aIF1 + GTP | 50S + aIF5B-H82A + GTP | 10.8 ± 1.2 |
| 8-30S:Met-tRNA<sub>i</sub><sup>Met</sup>:mRNA:aIF2:aIF1:aIF1A + GTP | 50S + aIF5B + GDPNP | 13.8 ± 1.8 |
| 9-30S:Met-tRNA<sub>i</sub><sup>Met</sup>:mRNA:aIF2:aIF1:aIF1A + GTP | 50S + aIF5B-ΔC + GTP | 3.8 ± 0.5 |

### Crystal structures

Crystals of full-length aIF5B (3 mg/mL, 0.5 mM GDPNP-Mg^2+^) were obtained at 4°C using 0.2 M Lithium nitrate and 20% PEG3350 (PEG Suite I, Qiagen) as precipitating agent. Crystals of aIF5B-ΔC (16 mg/mL, 10 mM GTP-Mg^2+^) were obtained in a solution containing 8% Tacsimate, 20% PEG3350 (PEG-ION 2 screen, Hampton Research). Before data collection, crystals of full-length aIF5B or aIF5B-ΔC were transferred in a solution containing the precipitating agent plus 25 % glycerol and then flash-cooled in liquid nitrogen. Diffraction data were collected at 100 K, λ= 0.98 Å, on the Proxima-1 (aIF5B) and Proxima-2 (aIF5B-ΔC) beamlines, at the SOLEIL synchrotron (Saint-Aubin, France).

Diffraction images were analyzed with XDS (62) and processed with programs of the CCP4 package (63). The structure of full-length aIF5B was solved by molecular replacement with PHASER (64) using the structure of aIF5B from *Aeropyrum pernix* (PDB ID, 5FG3, (39)) as a search model with 3 independent modules containing domains I and II, domain III and domain IV, respectively. Coordinates and associated B factors were refined through several cycles of manual adjustments with COOT (65) and positional refinement with BUSTER (66) and PHENIX (67). During the course of structure refinement, we noted the presence of a GDP molecule within the active site although the crystals were prepared in the presence of 0.5 mM GDPNP-Mg^2+^. Final statistics are shown in Supplementary Table 1. The final model contains all residues (from 2 to 598) with one GDP molecule bound to the active site.

No magnesium atom was visible in the electron density. The structure of aIF5B-ΔC:GTP was solved by molecular replacement using three independent structural domains corresponding to domain I, II and III of full-length aIF5B. The same strategy as that described above was used for refinement. The final model was refined to 1.7 Å resolution (Supplementary Table 1). It contains residues 3-455 bound to GTP, Mg^2+^ and Na^+^. Parts of domain III and of the h12 helix that do not interact with domains I and II show weaker electron density and higher B-values.

### IC3 complex preparation and cryo-EM analysis

The strategy used to prepare the 30S:mRNA:aIF1A:aIF5B:Met-tRNA_i_^Met^ complex, hereafter named IC3, was adapted from that used for other *P. abyssi* initiation complexes (19,20). First, archaeal 30S subunits from *P. abyssi* (Pa-30S) were heated 5 minutes at 51°C before mixing with a two-fold excess of Met-tRNA_i_^Met^, a five-fold excess of a synthetic 26 nucleotide-long mRNA (A_(−17)_UUUGGAGGUGAUUUAAA_(+1)_UGCCAAAG_(+9)_, ThermoScientific) in buffer A (10 mM MOPS pH 6.7, 100 mM NH_4_Cl, 10 mM magnesium acetate, 3 mM 2-mercaptoethanol). The mixture was then incubated 1 minute at 51°C. aIF5B was incubated 5 min at 51°C with 1 mM GDPNP-Mg^2+^. Then, a five-fold excess of initiation factors aIF5B:GDPNP and aIF1A was added to the 30S:mRNA:Met-tRNA_i_^Met^ complex. The IC3 complex was further incubated 2 min at 51°C and 5 min at room temperature. Finally, IC3 was purified by affinity chromatography (TALON resin, Clontech) using the two N-terminal histidine tags of aIF5B and aIF1A. IC3 was then dialyzed against buffer A to remove imidazole, concentrated using centricon 100K and stored at -80°C after flash-freezing into liquid nitrogen. The presence of all components was confirmed by SDS-PAGE and Western blot analysis (Supplementary Figure 1). Before spotting onto R2/1 grids with an extra 2 nm carbon layer (Quantifoil, Inc), the complex was diluted to a final concentration of ∼100 nM and a 5-fold excess of Met-tRNA_i_^Met^, aIF1A and aIF5B was added. For the final data collection, in order to stabilize aIF5B onto the SSU, we used BS^3^ ((bis(sulfosuccinimidyl)suberate), Thermo Fisher Scientific) as a crosslinker. Before crosslinking, purified IC3 was dialyzed against buffer B (10 mM MOPS pH 6.7, 100 mM NaCl, 10 mM magnesium acetate, 3 mM 2-mercaptoethanol) to remove ammonium that could react with BS^3^. The complex was diluted to a final concentration of ∼100 nM after addition of a 5-fold excess of Met-tRNA_i_^Met^, aIF1A and aIF5B. IC3 was then heated 1 minute at 51°C before adding BS^3^ at a final concentration of 1.5 mM and further incubated 10 to 15 minutes at 51°C. Immediately after, a 3.4 μL sample from the mixture was applied on the grid at 20°C and 90% humidity for 10 s. The sample was vitrified by plunging into liquid ethane at -182 °C, after 1.2 s blotting (with the sensor option) using a Leica EM-GP plunger. Cryo-EM images were collected on an FEI Titan Krios microscope (Thermo Fisher Scientific) operated at 300 kV at Institut Pasteur, equipped with a K3 direct electron detector (GATAN) at a pixel size of 0.86 Å/pixel. 40 frames were collected for 2.35 s in linear mode with defoci ranging from -0.8 μm to -3.0 μm, and an exposure rate of 17 e-/Å^2^/s. In total, 19.1 k micrographs were collected (Supplementary Table 2).

### Data processing

Motion correction was performed using RELION’s implementation (68). The frames were aligned using 5×5 patches with 1 e-/Å^2^/frame dose-weighting. CTF was estimated using gCTF (69) on the dose-weighted micrographs. Aligned micrographs with gross surface contamination and obvious gCTF output-metrics outliers were discarded. Power spectra with nominal resolutions better than 4 Å, objective-lens astigmatism better than 350 Å and within a [0.7 -3.8] μm defocus range were kept for further processing. All steps of data processing were performed in RELION 3.1 (68). The final pool comprised 17.1 k micrographs, 89% of the initial dataset. Particles were extracted using 432 × 432 pixel boxes. To reduce computation time of the initial stages of processing, particles were downscaled 3-fold to a 144×144 pixel box size (2.58 Å pixel size). The resulting final pool had 2.6 million particles with an average of 153 particles per micrograph. After several rounds of 2D-classification, 2 million particles remained. An initial 3D refinement with 2 million particles gave a 5.1 Å resolution density map (Supplementary Figure 2). Next, a 3D-classification job (8 classes, no sampling) isolated one main class showing all components of IC3, mRNA, initiator tRNA, aIF5B and aIF1A (1.1 M particles). The other classes contained only parts of IC3 with poorly defined 30S subunit and were not further used. The full-size re-extracted 1.1 M particles were further used to refine the 30S high-resolution cryo-EM map. After particle CTF refinement and post-processing, Map A (2.25 Å resolution) was obtained (Supplementary Table 2). A second round of 3D-classification of the 1.1 M binned particles with local sampling led to the identification of 4 classes with one class containing all components (Class 4, 118 k particles). The full-size re-extracted particles from Class 4 allowed us to refine a cryo-EM map at an overall resolution of 2.9 Å. In final steps of refinement, the quality of the cryo-EM map of IC3 (30S:mRNA:aIF1A:aIF5B:Met-tRNA_i_^Met^) was further improved by using masked 3D classification to account for conformational heterogeneity, CTF refinement and Bayesian polishing. This yielded a 2.7 Å resolution IC3 cryo-EM map calculated from 37 k particles (Map B). The map around aIF5B domain IV was still at lower resolution than for the rest of the initiation factor. Thus, we used multi-body refinement (70) to further improve the map quality of aIF5B and of the acceptor helix of Met-tRNA_i_^Met^ (Supplementary Figures 2 and 3). Two soft masks were created using RELION. The first mask covered the 30S subunit and the mRNA (Body 1) and the second one covered aIF1A:aIF5B:Met-tRNA_i_^Met^ (Body 2). These masks were used for multibody refinement in RELION, which yielded partial maps B1 and B2 with 2.6 Å and 3.6 Å overall resolution, respectively. Particles from Class 2 and Class 3 were also re-extracted at full resolution, refined and 3D classified to improve the densities of the factors and tRNA. Class 2 contained particles corresponding to 30S:mRNA:aIF1A:Met-tRNA_i_^Met^ and class 3 contained particles corresponding to 30S:mRNA:aIF1A:aIF5B-DIV:Met-tRNA_i_^Met^. The overall resolutions of the final maps obtained with Class 2 and 3 were 2.6 Å and 2.8 Å, respectively (Supplementary Figure 2). Class 1 only showed aIF1A and mRNA and was not further processed. The resolutions of all maps were obtained with RELION’s post-processing tool based on the gold-standard Fourier shell correlation (FSC) using the 0.143 cut-off criterion. The local resolution range was estimated using RELION’s local resolution tool (71).

### IC3 model building and refinement

The structure of the 30S was first manually adjusted in map A (2.25 Å resolution) using Coot (65) and *P. abyssi* 30S subunit (PDB ID 6SWC) as an initial model (19). In particular, a total of 67 rRNA modifications were modeled and the 16S rRNA structure was updated accordingly (Supplementary Table 3). The mRNA and the tRNA anticodon stem-loop (ASL) were also modeled in Map A (Supplementary Figure 4). Then, the complete 30S:mRNA:Met-tRNA_i_^Met^-ASL model was refined through cycles of manual adjustments in Coot (65) and energy minimization in Phenix (72). This improved model of 30S:mRNA:Met-tRNA_i_^Met^-ASL was used as a starting model for building IC3, 30S:mRNA:aIF1A:aIF5B-DIV:Met-tRNA_i_^Met^ and 30S:mRNA:aIF1A:Met-tRNA_i_^Met^ structures.

The models of 30S:mRNA:aIF1A:aIF5B-DIV:Met-tRNA_i_^Met^ and 30S:mRNA:aIF1A:Met-tRNA_i_^Met^ were built and refined in the corresponding unsharpened maps using Phenix (maps deposited as EMD-14579 and EMD-14581). aIF5B-DIV and aIF1A were poorly defined and the models of these two proteins were placed in the density without further positional refinement. The models of 30S:mRNA:aIF1A:aIF5B-DIV:Met-tRNA_i_^Met^ and 30S:mRNA:aIF1A:Met-tRNA_i_^Met^ were only used to analyze the position of the initiator tRNA. For IC3, models of aIF5B (this study, PDB IDs 7YYP and 7YZN), aIF1A ((19), PDB ID 4MNO) and Met-tRNA_i_^Met^ ((57), PDB ID 5L4O) were first adjusted in Map B. The sharpened multibody map B2 was then used to manually adjust and refine the model of aIF1A:aIF5B:Met-tRNA_i_^Met^ (Supplementary Table 2). To build the final IC3 model, we used the strategy described in (70). We refined the model of the 30S subunit into map B, placed the above model of aIF1A:aIF5B:Met-tRNA_i_^Met^and finally refined the B factors (Supplementary Table 2). Refinement statistics (73) are given in Supplementary Table 2. All figures were done with ChimeraX (74) or Pymol (75).

## RESULTS

### aIF5B hydrolyzes GTP on the ribosome and accelerates subunit joining

To test the activity of aIF5B, we first measured the rate of γ-[^32^P]GTP hydrolysis after mixing 30S complexes and 50S subunits (see Methods). In the presence of the 30S and 50S subunits only, aIF5B hydrolyzed GTP with a rate constant of 0.25 ± 0.02 min^-1^ (Table 1, line 4 and Figure 1A). The GTPase activity of aIF5B was dependent on the presence of both 30S and 50S (Table 1, lines 1-4). This result is consistent with previous studies on eIF5B from human (30,31) or *Kluyveromyces lactis* (43) and from *S. solfataricus* aIF5B (51). When Met-tRNA_i_^Met^ and mRNA were added to the 30S, GTP hydrolysis rate increased to 0.35 ± 0.04 min^-1^. The addition of aIF1A only marginally influenced this rate constant (0.32 ± 0.02 min^-1^, Figure 1A and Table 1, lines 5-6). These results show that aIF5B is active for GTP hydrolysis on the assembled ribosome and that this activity mainly reflects the binding of aIF5B close to the GTPase center on the LSU. Accordingly, GTPase activity was still observed when aIF5B was replaced by aIF5B-ΔC that does not bind Met-tRNA_i_^Met^ (Table 1, compare lines 9-10 to lines 6 and 4). We therefore hypothesized that GTP hydrolysis on aIF5B was triggered by the histidine residue (H82) of the switch 2 region conserved in all translational GTPases (Supplementary Figure 5A). Indeed, the role of the equivalent histidine in GTP hydrolysis was already evidenced using human (H706) or yeast (H480) eIF5B (28,31,33). Consistent with this idea, the aIF5B-H82A variant did not exhibit any detectable γ-[^32^P]GTP hydrolysis activity, showing that H82 is essential for the GTPase activity of aIF5B (Table 1, line 7 and Figure 1A).

We then studied the role of aIF5B in subunit joining. Association of ribosomal subunits was monitored with light scattering at 51°C after rapid mixing using a stopped-flow apparatus (see Methods) as previously described for bacterial and eukaryotic systems (32,60,61,76). Very slow joining was observed upon mixing 30S and 50S subunits in the absence of other components (Figure 1B, black curve, Table 2, line 1). Then, we prepared 30S initiation complexes by mixing 30S subunits, mRNA, Met-tRNA_i_^Met^, aIF1, aIF1A and aIF2 in the presence of GTP. Preliminary measurements showed that γ- [^32^P]GTP hydrolysis by aIF2 under these conditions occurred at a rate of 0.22 ± 0.03 min^-1^. Thus, the 30S initiation complexes were incubated at least 30 minutes at 51°C to allow full GTP hydrolysis, followed by aIF1 and aIF2:GDP departure, before mixing with 50S subunits in the presence of aIF5B and GTP. The scattering curve showed rapid association of the subunits (Figure 1B, blue curve). The data were fitted to hyperbolic curves as described (61) from which a rate constant of 11.4 ± 3.0 min^-1^ was derived (Table 2, line 5). As observed in the eukaryotic case, GTP hydrolysis was not required for the joining step since aIF5B could be replaced by its H82A variant (Figure 1B, orange curve) or GTP by GDPNP, without significant modification of the association rate (Table 2, lines 5,6,8). Conversely, in the presence of all components except aIF5B, subunits were unable to assemble (Figure 1B, brown curve and Table 2, line 4). However, under our assay conditions, we did not observe any significant effect of the presence of aIF1A on aIF5B-promoted subunit joining (Table 2, lines 6,7). On another hand, mRNA and Met-tRNA_i_^Met^ were sufficient to promote subunit association, though at a much slower rate (0.32 ± 0.04 min^- 1^; Table 2, line 3; Figure 1B, light grey curve). Similarly, aIF5B:GTP (H82A variant) alone also promoted subunit association (0.84 ± 0.06 min^-1^; Table 2, line 2; Figure 1B, dark grey curve). Consistent with a synergistic action of aIF5B and Met-tRNA_i_^Met^, removal of the C-terminal domain of aIF5B (aIF5B-ΔC), known to interact with the tRNA, lowered the association rate by a factor of 3 as compared to full-length aIF5B (Table 2, lines 5 and 9).

### Structure of full-length aIF5B bound to GDP

The overall crystallographic structure of full-length *P. abyssi* aIF5B shows the four domains arranged as described previously for other eukaryotic (36,37,40) or archaeal (38,39) e/aIF5B (Figure 2A). Consistent with the presence of GDP in the active site, the two switch regions involved in the binding of the nucleotide adopt the OFF conformation as previously observed in the two other aIF5B:GDP structures (*Aeropyrum pernix*, PDB ID 5FG3 (39) and *Methanobacterium thermoautotrophicum*, PDB ID 1G7S (38)) or in the apo eIF5B from *S. cerevisiae* (PDB ID 3WBI (40)). Domains I of all these structures can be superimposed with very small rmsd values (Supplementary Figure 6). According to this superimposition, position of domain II only weakly varies whereas position of domain III varies significantly in the different structures. Finally, domain IV appears to move freely with respect to domains I, II and III (Supplementary Figure 6).

**Figure 2:**
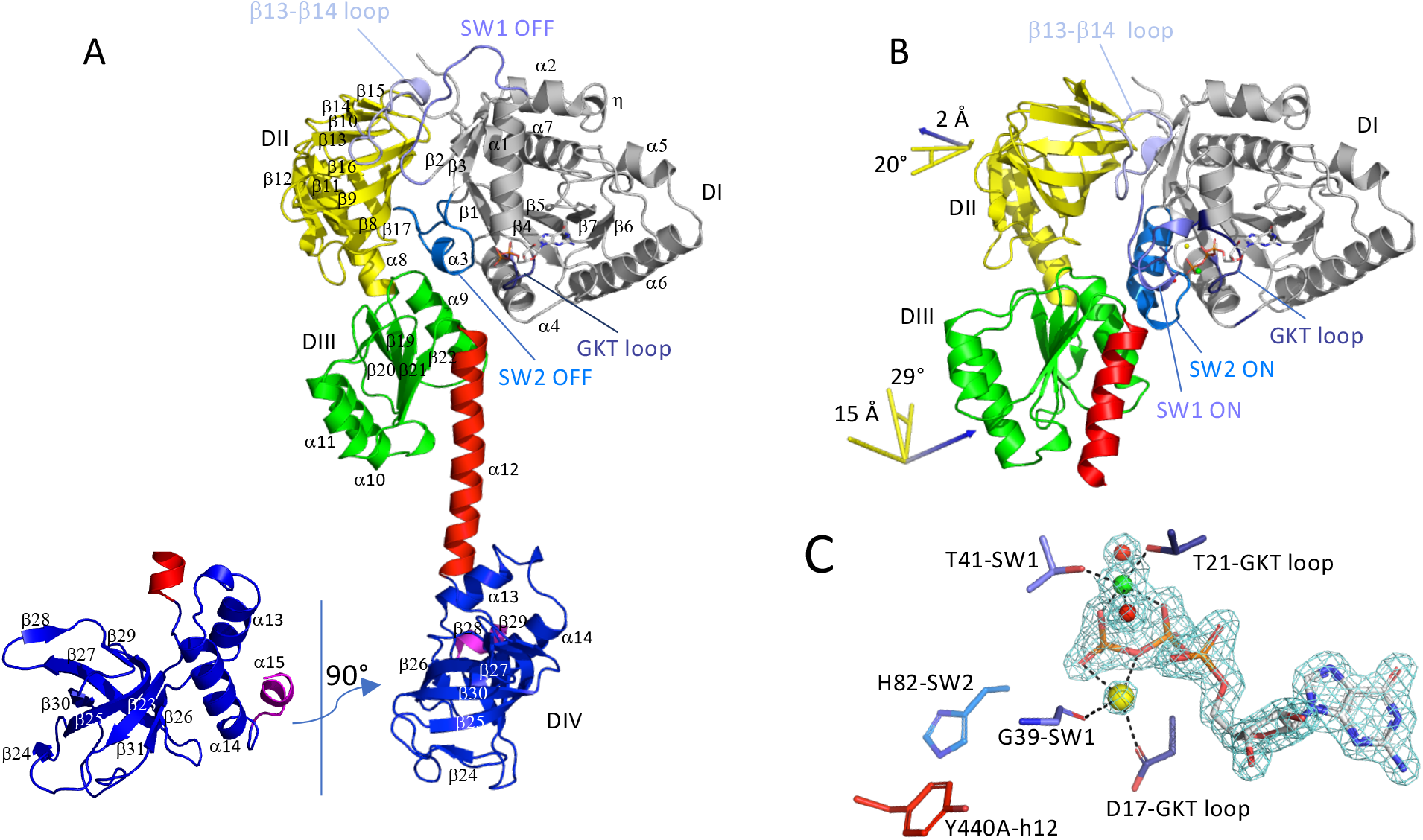
Crystallographic structures of aIF5B from *P. abyssi*. (**A**) Cartoon representation of full-length aIF5B. The aIF5B domains are colored as follows; domain I gray (1-229), domain II yellow (230-348), domain III green (350-438), α12 red, (439-464), domain IV blue (465-591) and the C-terminal α15 helix magenta (591-598). The GDP is shown as sticks and main regions involved in the binding of the nucleotide are colored as follows, GKT loop deep blue, switch 1 (SW1) slate blue, switch 2 (SW2) marine and the β13-β14 loop of domain II light blue. Domain IV is shown in two orientations. (**B**) Cartoon representation of aIF5B-ΔC. The color code is the same as in view A. Domains I of the two structures have been superimposed. Movements of domains II and III from aIF5B to aIF5B-ΔC have been calculated with Pymol. Rotation axes are shown as blue arrows, rotation angles and translations are indicated beside domains II and III. (**C**) GTP binding site. The 1.7 Å resolution ‘2Fo-Fc’ map contoured at 2.6 standard deviation is drawn using the carve command of Pymol. GTP, magnesium (green), water (red) and the sodium (yellow) with important residues of SW1, SW2, the GKT loop and h12 are shown.

### Conformational changes associated with GTP binding

Prior to this study, the GTP-bound state of e/aIF5B outside of the ribosome with the switch ON conformation was only observed in a truncated version of eIF5B from *Chaetomium thermophilum* containing only domains I and II (Ct-DI-DII-eIF5B-GTP (36,37)). However, on the 80S ribosome, the GTP-bound form of the full-length factor was observed (42,43,45). These structures showed that domain III of eIF5B was involved in the stabilization of the active GTP-bound conformation.

We designed a *P. abyssi* aIF5B variant in which the C-terminal domain was deleted. The resulting protein (called aIF5B-ΔC) comprises domains I to III and the N-terminal part of the α12 helix (residues 1 to 458). As shown in Figure 1A, aIF5B showed no intrinsic GTPase activity in the absence of the ribosome. Consistent with this result, we could determine the crystal structure of aIF5B-ΔC bound to GTP-Mg^2+^ at 1.7 Å resolution (Supplementary Table 1). GTP binding involves the conserved sequences forming the nucleotide binding pocket of all G proteins (Figure 2B-C and Supplementary Figures 5). One magnesium and one sodium ion participate in GTP stabilization, as previously observed in *C. thermophilum* eIF5B-(DI-DII) truncated form (36,37). Interestingly, two metal ions were also observed in aIF2-GTP (77) showing that a second metallic atom participates in GTP binding in several translational GTPases. The two switch regions are ON, contributing to the binding of the γ-phosphate group of GTP. Additionally, the ON conformation is stabilized by a long loop of domain II located between β13 and β14 (residues 285 to 297, Figure 2B). Actually, the ON-OFF transition of the two switch regions is accompanied by a large movement of the β13-β14 loop of domain II (Figure 2). Such a movement was also noted in *C. thermophilum* eIF5B ((37) and Supplementary Figure 7A). Thus, the concerted rearrangement of switch 1 and switch 2 and of the β13-β14 loop during the GTP cycle of e/aIF5B is conserved in archaea and eukaryotes. Close to the GTP binding site, the side chain of H82, the universal histidine residue responsible for GTP hydrolysis (as shown here for aIF5B, Table 1), is in its non-activated conformation (Figure 2C). Moreover, at the N-terminal extremity of h12 is the strictly conserved residue Y440 (Figure 2C and supplementary Figure 5). This tyrosine was previously proposed to be important for GTP hydrolysis on eukaryotic eIF5B (42,43). However, no significant effect on GTPase activity was observed in our assay upon Y440A modification (Table 1, line 8) and further studies are required to fully understand the role of Y440 in *P. abyssi* aIF5B.

On another hand, the transition between the GDP and GTP states of aIF5B is accompanied by a movement of domains II and III illustrated in Figure 2. Notably, the position of domain II with respect to domain I in *P. abyssi* aIF5B-ΔC:GTP is very similar to that observed in Ct-DI-DII-eIF5B:GTP (37) (Supplementary Figure 7A and B). Globally, the arrangement of domains I, II and III in aIF5BΔC:GTP is close to that observed in eukaryotic eIF5B bound to the 80S ribosome. This point will be discussed more deeply below. Of note, weaker quality of the electron density and higher B-values were observed in parts of aIF5B-ΔC domain III and of the h12 helix that do not interact with domains I and II. Interestingly, the corresponding regions of *S. cerevisiae* eIF5B interact with the ribosome in eIF5B:80S complexes (42,43,45,46). Finally, the present crystallographic structure shows that aIF5B does not require the SSU or the ribosome as a cofactor to bind GTP and trigger domains I, II and III rearrangement.

### Overview of the IC3 cryo-EM structure

To analyze the molecular events occurring on the SSU after aIF2 departure and before its joining with the large ribosomal subunit, we prepared an initiation complex containing the small ribosomal subunit bound to a synthetic 26 nucleotide RNA derived from *P. abyssi* aEF1A mRNA, containing a strong SD sequence (20), Met-tRNA_i_^Met^, aIF5B:GDPNP and aIF1A (see Materials and Methods). This complex, hereafter called IC3, was further purified by affinity chromatography using N-terminally His-tagged versions of aIF5B and aIF1A (Supplementary Figure 1). An excess of Met-tRNA_i_^Met^, aIF1A, aIF5B was added before spotting IC3 onto the grids for cryo-EM data collections. After image processing, although the mRNA, the tRNA and aIF1A were clearly visible, we systematically only observed a density blob bound to the methionylated tRNA acceptor end, likely corresponding to domain IV of aIF5B but density corresponding to domains I-II-III of aIF5B on the 30S was only faintly visible (data not shown). This suggested that aIF5B binding to the 30S was not stable enough in our experimental conditions. Therefore, in order to improve the quality of the cryo-EM maps, we crosslinked IC3 using bis(sulfosuccinimidyl)suberate (BS^3^) at 51°C as previously done for other ICs (*e*.*g*. (78,79), see Materials and Methods). Cross-linking with BS^3^ did not change the overall structure, but it greatly improved the quality of the potential map in the aIF5B binding regions.

Cryo-EM images were collected on a Titan Krios microscope (Supplementary Table 2) yielding a total of 19.1 K micrographs. Images were processed using RELION 3.1 (68,80,81) (Supplementary Figure 2; Materials and Methods). Briefly, an initial dataset with 2 M particles was used for an initial 3D-reconstruction in which all components of IC3 were visible. Several rounds of 3D-classification identified four main classes. Class 4 contained 118 k particles with all factors visible. Class 3 contained 522 k particles with 30S, mRNA, aIF1A, tRNA but only domain IV of aIF5B was visible. Class 2 (400 k particles) contained 30S bound to tRNA, mRNA and 1A. Finally, Class 1 (131 k particles) contains 30S bound to mRNA and 1A only. After a round of masked-3D classification to account for conformational heterogeneity, we finally isolated from Class 4 a subset of 37 k particles that yielded a potential map at 2.7 Å resolution (map B). In a final processing step, we used a multibody refinement strategy to improve the potential map in the aIF5B:tRNA:aIF1A region. Particles from Classes 2 and 3 were further 3D classified to improve densities of the factors and tRNA. Using gold-standard FSC curves, the final resolution of maps from Classes 2 (30S:aIF1A:Met-tRNA_i_^Met^), 3 (30S:aIF1A:aIF5B-DIV:Met-tRNA_i_^Met^) and 4 (IC3) were 2.6 Å, 2.8 Å and 2.7 Å, respectively.

Map B clearly showed well-defined density for all IC3 actors. Thanks to the multibody refinement strategy, we obtained map B2 where the Met-initiator tRNA, aIF1A and aIF5B shows improved local resolutions ranging from 2.4 to 6 Å (Figure 3 A,B and Supplementary Figure 3). Overall, aIF5B is located on the SSU at the expected position as deduced from the cryo-EM structures of the eukaryotic 80S complexes containing eIF5B (34,42,46,82).

As shown in Figure 4A, the initiator tRNA is fully base paired with the AUG start codon in the P site and the position of the ASL (anticodon stem-loop) is stabilized by numerous interactions as previously observed in the 30S:mRNA:Met-tRNA_i_^Met^:aIF2:aIF1A (called IC2) complex illustrating the preceding step of TI (19). Among them, R135 of uS9 interacts with the phosphate groups of U33, C34 and A35 in the anticodon loop (Figure 4A). In the C-terminal tail of uS19, T123 and S125 interact with the phosphoryl of G30 and uS19-R124 is stacked along the phosphate backbone of G30. The NH group of uS13-V145 interacts with the phosphoryl of G29 (Supplementary Figure 8). The P gate nucleotides, G1312 and A1313, interact with the G29-C41 and the G30-C40 base pairs of the anticodon stem. On Sthe other side, the pocket is delineated by A757 from h24 loop. aIF1A is stably bound in the A site and in the exit channel, the mRNA is base-paired to 9 bases of the 3’ extremity of the 16S rRNA, as observed previously (19).

At the tRNA extremity, the 3’ methionylated moiety is bound to aIF5B-domain IV. aIF5B-domain IV also contacts aIF1A (Figure 3D) bound to the A site. Helix h12 of aIF5B is well defined, connecting domain IV to the three-domain core (domains I, II and III) of aIF5B. Domains II and III interact with the shoulder and the body regions of the 30S subunit (Figure 3C). In domain I, the switch regions are in the “ON” conformation and GDPNP-Mg^2+^ is clearly visible (Figure 4B). H82 of switch 2 and Y440 at the N-ter extremity of h12 have the same conformations as in aIF5B-ΔC:GTP structure.

**Figure 3:**
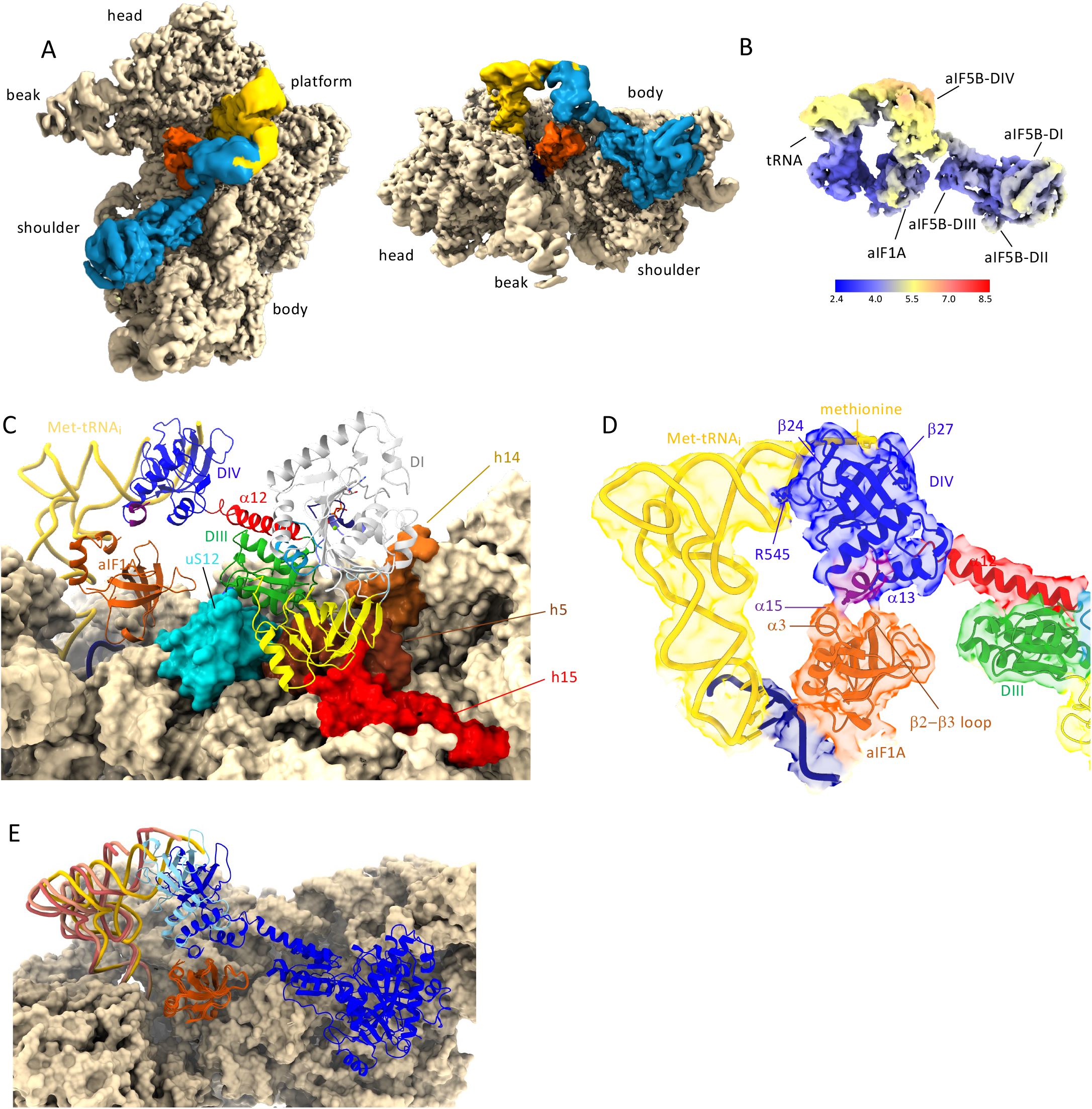
Cryo-EM structure of IC3. (**A**) Map B colored according to its composition, the SSU in beige, Met:tRNA_i_^Met^ in bright yellow, aIF1A in orange and aIF5B in blue. The map is shown in two orientations. (**B**) Local resolution of the multi-body refinement map B2 calculated using the script implemented in RELION (71). (**C**) IC3 structure showing interaction of aIF5B with the SSU and aIF1A.The color code is the same as in Figure 2. uS12 is in cyan. h15 is in red, h5 and h14 are in brown. The mRNA is in dark blue. (**D**) Closeup showing the cryo-EM map obtained after multibody refinement around aIF1A and aIF5B. The color code is the same as in Figure 2. (**E**) Superimposition of 30S:aIF1A:Met-tRNA_i_^Met^ and 30S:aIF1A:aIF5B-DIV:Met-tRNA_i_^Met^ subcomplexes onto IC3 showing the positions of the initiator tRNA in the three structures. Met-tRNA ^Met^ is dark pink in 30S:aIF1A:Met-tRNA_i_^Met^, light pink in 30S:aIF1A:aIF5B-DIV:Met-tRNA_i_^Met^ and yellow in IC3. Domain IV of aIF5B is light blue in 30S:aIF1A:aIF5B-DIV:Met-tRNA_i_^Met^ and dark blue in IC3. This view shows that the conformation of the initiator tRNA is constrained by the interactions of aIF5B with the SSU and with aIF1A.

**Figure 4:**
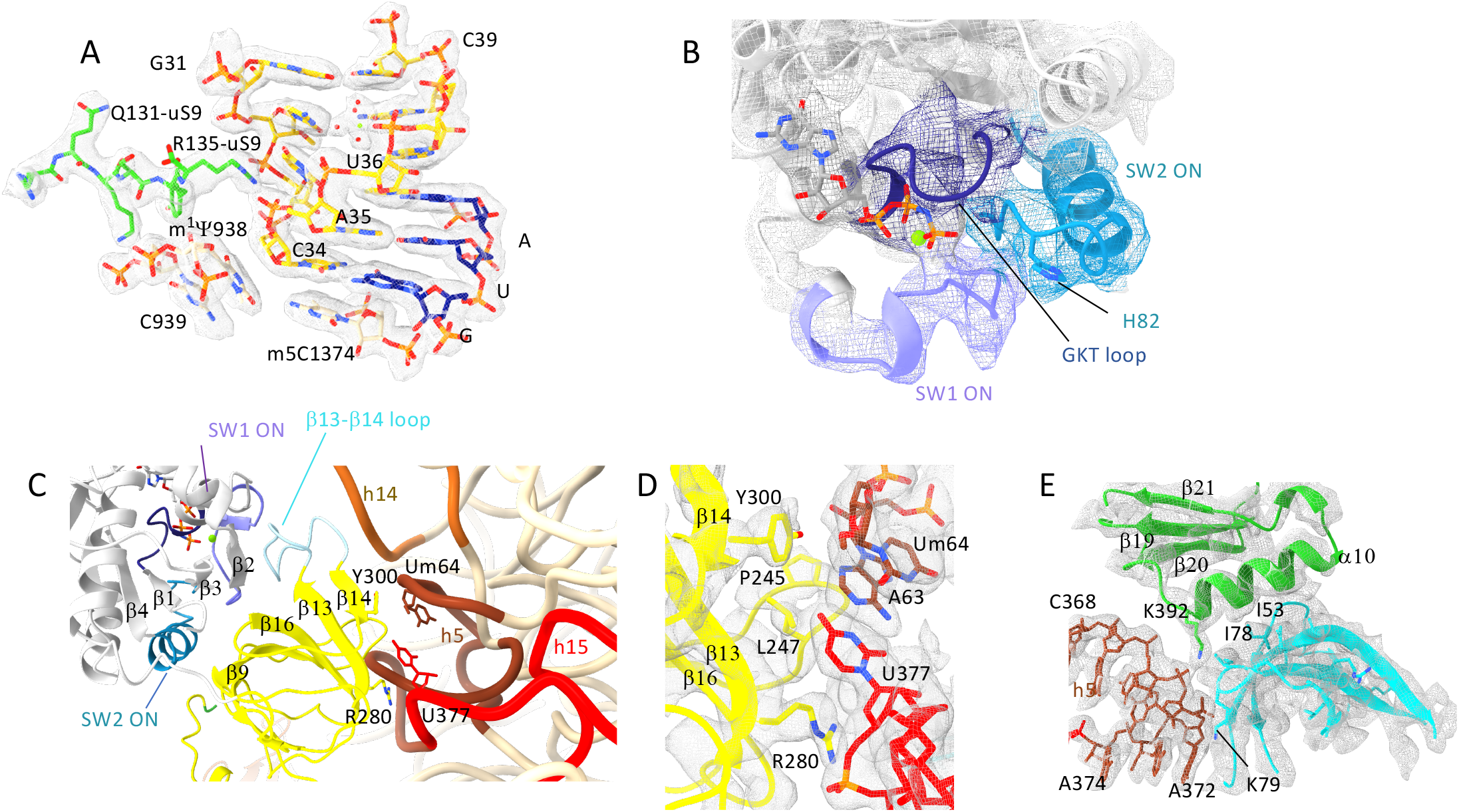
Met-tRNA_i_^Met^ and aIF5B in IC3. (**A**) Closeup of map B showing the codon:anticodon interaction at the P site. The C-terminal tail of uS9 is shown in green. mRNA bases are in blue, rRNA in beige and tRNA in yellow. Another orientation is shown in Supplementary Figure 8. (**B**) Closeup of map B2 around GDPNP. The color code is the same as in Figure 2. (**C**) Interaction of aIF5B-DII with the 30S. See also the text and Supplementary Table 4 for the description of the important residues shown here as sticks. (**D**) Closeup of map B2 at domain II and SSU interface. (**E**)Closeup of map B2 at the interface of aIF5B-DIII and uS12.

### Interaction of aIF5B domain IV with Met-tRNA_i_^Met^ and aIF1A

Domain IV of aIF5B interacts with both aIF1A and the methionylated _73_ACCA_76_ end of the initiator-tRNA. The quality of map B2 was sufficient to position unambiguously the secondary structures and to highlight the role of several side chains (Figure 3D and Supplementary Figure 9). The methionylated end of the initiator tRNA is bound to domain IV within a crevice delineated by the β27- β28 turn on the one side and by β24 and the preceding loop on the other side. β25 and β30 form the floor of the crevice and provide residues involved in hydrophobic interactions with the methionyl group (Supplementary Figure 9 and Supplementary Table 4). Overall, the crevice is rather closed thereby protecting the methionine ester bond from hydrolysis. R483 is stacked on A1 of the tRNA and R482 is stacked on C74. Interactions also involve the β30-β31 loop with R545, strictly conserved in e/aIF5B (Supplementary Table 4), that contacts base G70 of Met-tRNA_i_^Met^in the major groove of the acceptor stem. The terminal adenosine is bound on the side of β27 and β30.

The C-terminal region of domain IV of aIF5B is positioned on the solvent side of aIF1A (Figure 3D). Map B2 unambiguously shows contacts between aIF5B-domain IV and aIF1A (Figure 3D). The C-terminal extremity of aIF5B (residues 590 to 598) containing the archaeal specific α15 helix (39) is packed onto a shallow groove delineated by the short C-terminal α3 helix of aIF1A on the one side and by the β2-β3 loop on the other side (sequence _41_CEDGKI_46,_ with C and D strictly conserved in archaea).

The N-terminal part of α13 in aIF5B also contacts aIF1A at the level of the β2-β3 loop. The surface of interaction between the two proteins is rather small (329 Å^2^, as calculated with Pymol (75)) and few polar contacts are present. Although the resolution of maps B and B2 in this region was not sufficient to accurately position all side chains, the model shows hydrophobic contacts involving aIF5B-P593 and aIF1A-E103. The side chain of aIF5B-R559 likely interacts with the main chain carbonyl group of aIF1A-G44.

The position of the initiator tRNA observed in IC3 was compared to those in the 30S:aIF1A:Met-tRNA_i_^Met^ and in the 30S:aIF1A:aIF5B-DIV:Met_i_^Met^-tRNA subcomplexes. No significant movement of aIF1A is observed in the three structures. In 30S:aIF1A:Met-tRNA_i_^Met^, the initiator tRNA is base-paired with the mRNA and the anticodon loop adopts a position similar to that in IC3. However, the rest of the tRNA adopts a more upright position (Figure 3E). When DIV of aIF5B is bound to the acceptor end of the initiator tRNA while domains I-II-III of 5B are not stably bound to the SSU, as observed in the 30S:aIF1A:aIF5B-DIV:Met-tRNA_i_^Met^ structure, the position of the initiator tRNA appears as intermediate between that in IC3 and that in 30S:aIF1A:tRNA. In the latter structure, aIF5B-DIV does not contact aIF1A. Therefore, the conformation of the initiator tRNA is constrained by the interaction of aIF5B with the SSU and with aIF1A. Overall, aIF5B maintains the Met-tRNA_i_^Met^ in a conformation adequate for 50S subunit joining, as discussed below.

### Interaction of aIF5B with the small ribosomal subunit

Apart from domain IV, the remaining of aIF5B interacts with the body of the SSU via its domains II and III. The surface of interaction is ∼1900 Å^2^. There is no interaction between domain I and the SSU. Three loops of domain II contact the SSU at the level of h15 in the shoulder and h5 in the body (Figure 4C,D and Supplementary Table 3). The β13-β14 loop is close to h14 but does not directly contact it. A notable interaction involves Y300 of the β13-β14 loop and 2’O-methyl U64 of h5 (Figure 4D). Y300 is however conserved neither in archaea nor in Thermococcales and its role cannot therefore be generalized. As also observed in the aIF5B-ΔC:GTP crystallographic structure, the β13-β14 loop of aIF5B-domain II interacts with switch 1, confirming that the position of this loop is correlated with the nucleotide state of aIF5B. Domain III contacts the h5-h14 junction and uS12 (Supplementary Table 3). uS12 is also in contact with aIF1A. The conformation of aIF5B bound to the SSU differs from that of unbound aIF5B:GDP (Figure 5A-B). In particular, the orientation of domain IV with respect to domain I has changed to ensure the binding of this domain with the initiator tRNA. This large movement of domain IV originates from the N-terminal extremity of the barrel. It is accompanied by a bending of helix h12 (Supplementary Figure 10).

**Figure 5:**
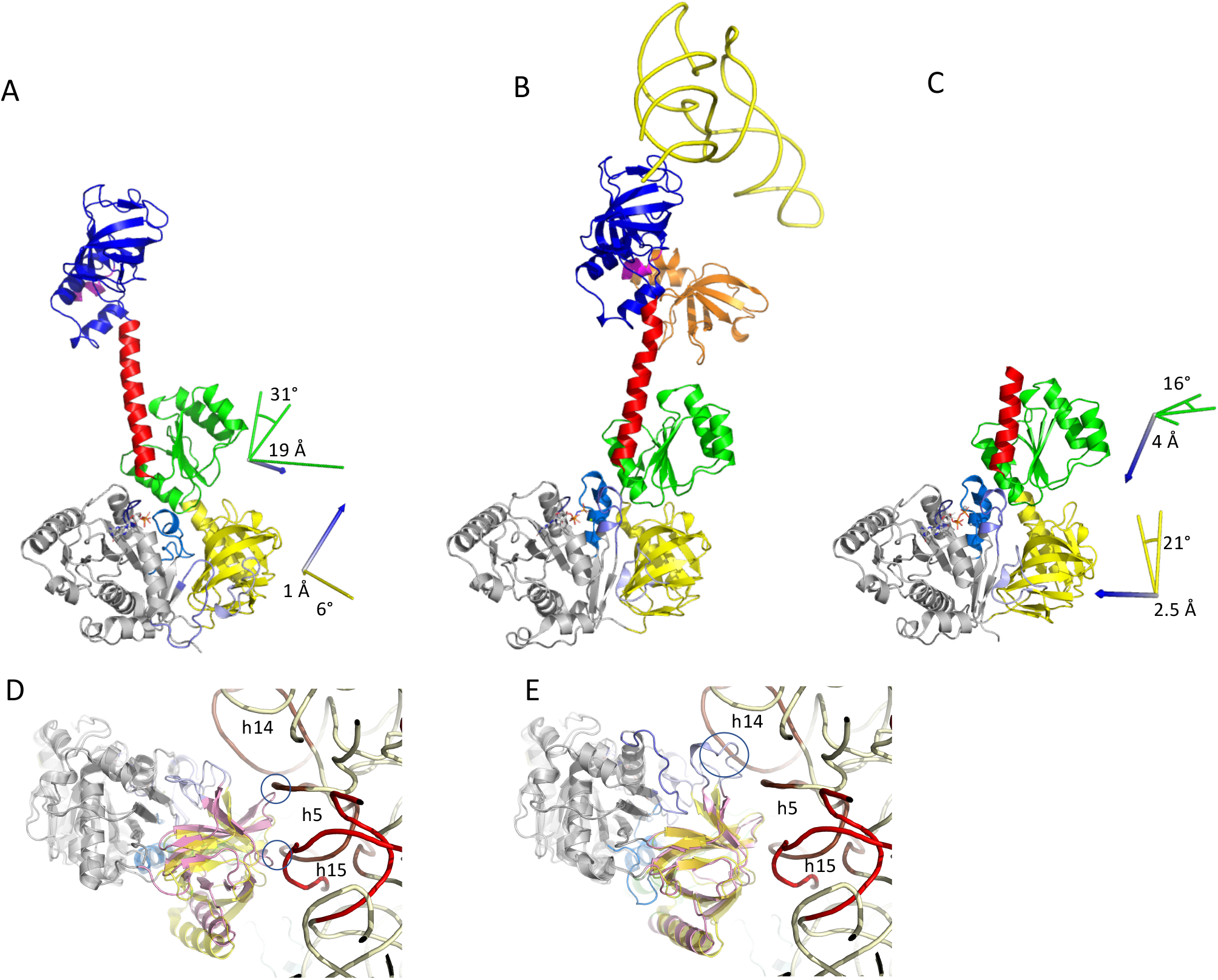
Comparison of Pa-aIF5B structures. In views (**A**), aIF5B:GDP, (**B**), aIF5B in IC3 and (**C**), aIF5B-ΔC:GTP, domains I of all structures were superimposed. Movements of domain II and III with respect of domain I were calculated with Pymol considering the structure of aIF5B in IC3 (B) as a reference. Movement of domain IV between aIF5B:GDP and aIF5B in IC3 calculated after superimposition of domains III is 35° rotation and 13 Å translation. (**D**) Domains I of aIF5B-ΔC:GTP and aIF5B in IC3 were superimposed. The color code is the same as in Figure 2 except that aIF5B-ΔC:GTP domain II is in pink. aIF5B-ΔC:GTP is shown at the foreground and aIF5B in IC3 is shown using transparent cartoons as a reference. The view shows that the position of domain II observed in aIF5B-ΔC:GTP would create steric clashes with h5 and h15. (**E**) Domains I of aIF5B:GDP and aIF5B in IC3 were superimposed. The color code is the same as in Figure 2 except that aIF5B:GDP domain II is in pink. aIF5B:GDP is shown at the foreground and aIF5B in IC3 is shown using transparent cartoons as a reference. The view shows the b13-b14 loop in the GDP conformation would create bad contacts with h14.

Interestingly, comparison of aIF5B:GDPNP bound to the SSU with the crystal structure aIF5B-ΔC:GTP shows some differences at the level of domains I, II and III orientations (Figure 5B,C). In particular, the position of domain II observed in aIF5B-ΔC:GTP would create bad contacts with h5 of rRNA (Figure 5D). Possibly, the initial binding of aIF5B to the SSU triggers a final adjustment of domain orientations in aIF5B leading to an optimal conformation of the factor for its binding to the GTPase activating center and the activation of subunit joining. In the same view, it is also notable that the position of the β13-β14 DII loop in the GDP state would strongly destabilize aIF5B binding on the SSU, suggesting a role of the rearrangement of this region in the release of aIF5B after GTP hydrolysis (Figure 5E).

### Comparison of the binding of the initiator tRNA to aIF2 with its binding to aIF5B

We previously determined cryo-EM structures of the first steps of TI in *P. abyssi* (19,20). After start codon recognition, the small initiation factor aIF1 is released from the ribosome and the initiator tRNA is fully accommodated in the P site of the SSU. The conformation of this complex was visualized in an IC2 cryo-EM structure containing the SSU, the mRNA, aIF1A and the ternary complex aIF2:GDPNP:met-initiator tRNA (19). In IC2, Met-tRNA_i_^Met^ is stably bound to the P site, aIF2 is released from its h44 binding site but still bound to the Met-tRNA_i_^Met^ because the non-hydrolysable GTP analogue, GDPNP, was used in the complex preparation. IC2 is representative of the initiation step just prior to aIF5B binding. To better understand conformational changes from IC2 to IC3, we superimposed rRNAs of the two structures (Figure 6A). A very small rsmd value of 0.6 Å was obtained for 30448 atoms compared. The overlay shows that the two structures are highly similar at the level of their rRNA, and that no movement of the SSU head occurs between the two initiation states. Moreover, the ASLs of Met-tRNA_i_^Met^ are very close (rotation angle 1.13° and 0.3 Å displacement) but the rest of the tRNA molecules differ (Figure 6A). A 13° rotation and a displacement of 5.6 Å between the upper parts of the tRNAs are measured (Figure 6A). This shows that this part of the tRNA bends towards the A site and the 30S body. The _73_ACCA_76_ methionylated end adopts a relaxed conformation when bound to domain IV of aIF5B compared to the IC2 conformation. Notably, aIF2γ and aIF5B approach Met-tRNA_i_^Met^ on opposite sides. This would allow a channeling of the Met-tRNA_i_^Met^ acceptor end and facilitate the transition between aIF2 departure and binding of aIF5B to the tRNA while minimizing exposure of the methionyl ester bond to the solvent.

**Figure 6:**
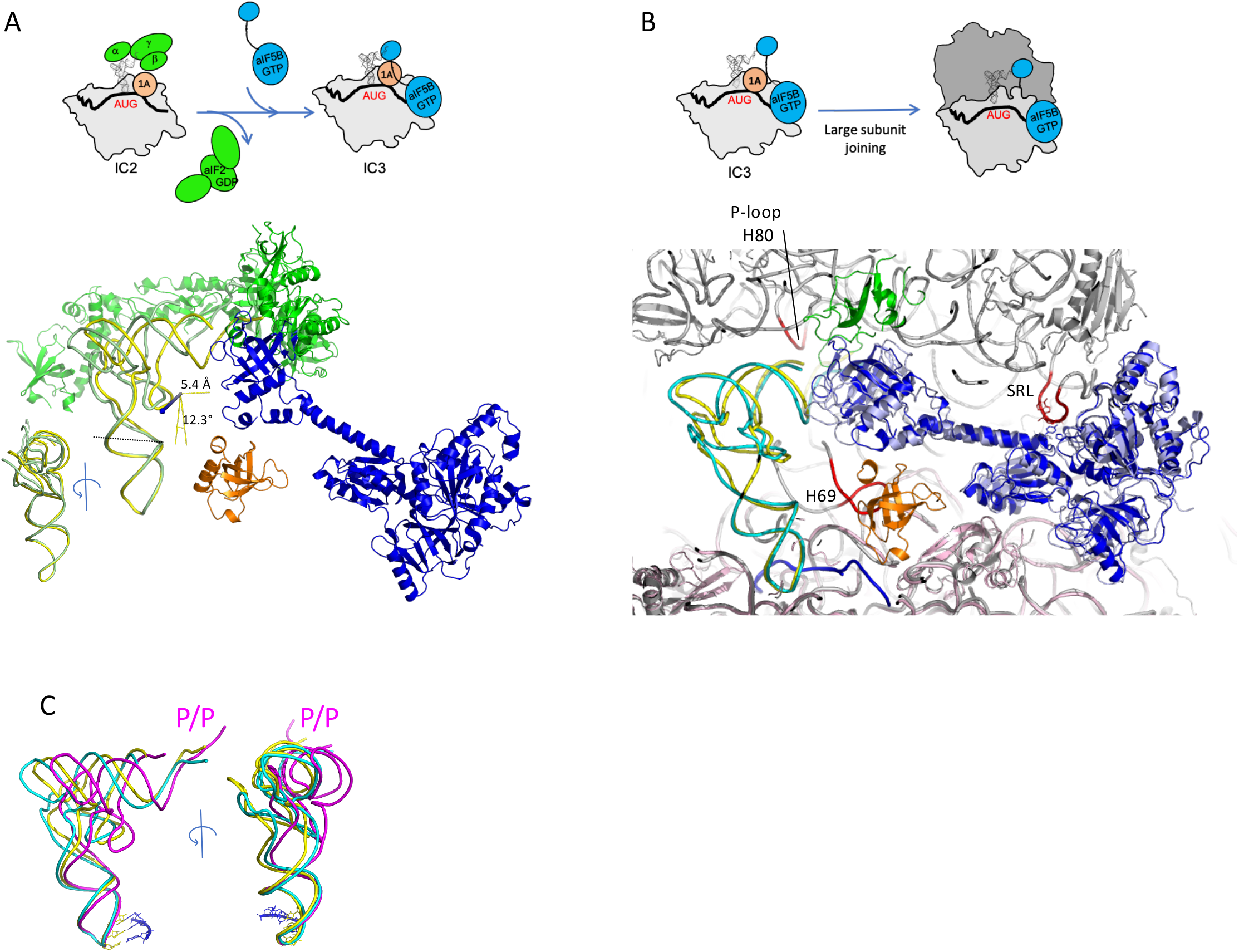
IC3 and the surrounding steps of translation initiation. (**A**) Comparison of IC3 with 30S:mRNA:aIF1A:aIF2:GDPNP:Met-tRNA_i_^Met^ (IC2). rRNAs of the the SSUs were superimposed (rsmd 0.6 Å for 30448 atoms compared). IC2 (PDB ID 6SWC, (19)) is colored as follows, aIF2α and γ subunits in green, tRNA in light green. IC3 is colored as follows; Met-initiator tRNA in yellow, aIF5B in blue, aIF1A in orange. The two anticodon stem-loops of the initiator tRNAs are very close (rotation 1.1° and 0.3 Å translation) but the rest of the tRNA molecules (nucleotides 1-26 and 44-76) differs by 13° rotation and 5.6 Å translation. The IC2 and IC3 steps are schematized on the left. (**B**) Comparison of IC3 with *S. cerevisiae* 80S:eIF5B complex. rRNA of the SSUs of *S. cerevisiae* 80S:eIF5B (PDB ID 6WOO, (46)) and IC3 were superimposed (see text). *S. cerevisiae* 80S:eIF5B is colored as follows; Met-initiator tRNA in cyan, aIF5B in light blue, uL16 in green. The P-loop of H80, H69 loop and SRL are colored in red. The ribosome is in grey. IC3 is colored as follows; Met-tRNA_i_^Met^ in yellow, aIF5B in blue, aIF1A in orange, 30S in light pink, Y440 and H82 are shown in blue sticks and A3029 of SRL is in red sticks. The ASLs of the two initiator tRNAs (nucleotides 27-43, rmsd=0.3; 1.9° rotation and 1.0 Å translation,) and domains I, II and III of e/aIF5B are very close (rsmd=1.14; 3.3° rotation and 1.0 Å translation). A small displacement between the upper parts of the two tRNAs (nucleotides 1-26 and 41-76, ∼5.7° rotation and 4.2 Å translation) with a concomitant movement of domain IV of e/aIF5B is observed (6.1° rotation and 4.6 Å translation). The view shows that the conformation of aIF5B:Met-tRNA_i_^Met^ in IC3 is almost equivalent to that of eIF5B:Met-tRNA_i_^Met^ in the *S. cerevisiae* 80S:eIF5B. Note that a positioning of H69 as observed in 6WOO would not be compatible with aIF1A. The IC3 model has been resized to allow superimposition onto 6WOO in order to take into account differences due to pixel size determination. (**C**) Binding state of the Met-tRNA_i_^Met^. Comparison of the Met-tRNA_i_^Met^ as observed in 6WOO (cyan) and IC3 (yellow) with the P/P tRNA as observed in a ribosome:EF-Tu:tRNA complex (PDB ID 5AFI, magenta, (97)).

### Pa-aIF5B and the joining step

We then used the IC3 structure to get insights into the joining step where aIF5B bound to the SSU becomes bound to the full ribosome. No structure of the archaeal 70S:aIF5B complex has been determined so far. Therefore, the present 30S:aIF5B structure was superimposed onto the structure of the *S. cerevisiae* 80S:eIF5B complex ((46), Figure 6B).

rRNAs of the small ribosomal subunits of the yeast 80S:eIF5B complex and of IC3 were superimposed with an rmsd value of 1.5 Å for ∼19,000 atoms compared. The overlay shows that the two SSUs are very similar without any head movement. Also, the two ASLs of the initiator tRNAs and domains I, II and III of e/aIF5B are very close (Figure 6B). Only a small displacement of the upper part of the tRNA with a concomitant movement of domain IV of e/aIF5B is observed (Figure 6B). Therefore, IC3 harbors an aIF5B:Met-tRNA_i_^Met^ conformation almost equivalent to that of eIF5B:Met-tRNA_i_^Met^ in the *S. cerevisiae* 80S:eIF5B complex. Only a small lowering of aIF5B-Domain IV and of the upper part of the Met-tRNA_i_^Met^would be necessary to stabilize the CCA-methionylated end close to the P-loop of the 23S rRNA of the LSU. As in eukaryotes, the aIF5B:Met-tRNA_i_^Met^ complex stabilization would involve the long extended loop of uL16. According to this superimposition, H82 of aIF5B would be perfectly placed with respect to the SRL loop that could trigger its active conformation and GTP hydrolysis, as observed for EF1A during the elongation process (83). In contrast, Y440 would create bad contacts with the SRL and should move upon 50S subunit joining towards a position as observed in yeast (46). Finally, the conformation of the *S. cerevisiae* 80S:eIF5B complex shows a positioning of the H69 helix that is not compatible with the position of aIF1A observed in IC3. This suggests that release of aIF1A would occur at an earlier step before H69 is completely placed close to the A site.

## DISCUSSION

The present study provides a structural and functional overview of the role of aIF5B when bound to the SSU. aIF5B:GDPNP is bound to the SSU through its domains II and III. Its domain IV is bound to the initiator tRNA and aIF1A. These interactions maintain the initiator tRNA in a conformation adequate for subunit joining.

Positioning of the methionyl moiety within the domain IV crevice is significantly different in *P. abyssi* as compared to *S. cerevisiae* (Supplementary Figure 9). The *P. abyssi* position would not be possible in the yeast system because of the presence of three inserted residues in the loop connecting β29 to β30. Such an insertion is not a general property of eukaryotes but is found in many fungi. Another difference between the Met crevice in *P. abyssi* aIF5B as compared to the yeast factor is the presence of a four-residue insertion between β23 and β24 (Supplementary Figure 9). This insertion, specific of Thermococcales, results in a more closed methionine site, likely contributing to better protect the ester bond in these hyperthermophilic organisms. Other archaea, and also many eukaryotes, have a two-residue insertion instead (Supplementary Figure 9). Therefore, whether the Met positioning in other eukaryotic aIF5B-DIV resembles that in *P. abyssi* or that in *S. cerevisiae* remains open. Overall, the Met binding pocket displays some variability that reflects specific adaptations. In the same view, recognition of the formyl-Met moiety involves specific bacterial features in domain IV of IF2 (84,85) (Supplementary Figure 9).

A long-lasting question concerns the interaction of factors 1A and 5B when bound to the small subunit of the ribosome before the assembly with the large subunit. This question is even more burning for the archaeal case than for the eukaryotic one because no biochemical data in favor of such an interaction is available. Indeed, aIF1A does not possess the eukaryotic-specific C-terminal eIF1A extension shown to mediate its interaction with eIF5B (35,40,48-50). In this study, we observe that aIF1A interacts with aIF5B on the SSU. This interaction involves the archaeal-specific C-terminal α15 helix as previously proposed (39). Although not conserved in sequence, α15 helix is almost systematically present in all archaeal aIF5B (Legend of Supplementary Table 4). In the absence of aIF5B, the initiator tRNA adopts a more upright position. Therefore, the interaction between aIF5B and aIF1A contribute to the positioning of the initiator tRNA and to the stabilization of the initiation complex in a conformation adequate for LSU joining (Figure 3E and 6B). This conformation would allow a facilitated accommodation of the CCA end in the peptidyl transferase center. In our study of the archaeal system, the light scattering monitored association assay did not show a significant lowering of the association rate when aF1A was omitted. Because the surface interaction between aIF1A and aIF5B is rather small (329 Å^2^), the effect of aIF1A might be difficult to evidence in a global association assay. A more detailed kinetic study, out of the scope of the present work, using bulk or single molecule FRET might help to better understand the dynamic interplay between the two factors.

Interactions between human eIF5B and eIF1A contributing to an adequate positioning of the initiator tRNA on the SSU were also observed in two eukaryotic studies published in BioRxiv (53,54) while this manuscript was being written. These interactions involve the two terminal α-helices of eIF5B and the 2-3 loop of eIF1A and contribute to a relatively small interaction surface (∼150 Å^2^). A similar contact is also observed in the present study. Surprisingly, interaction with human eIF5B do not involve the eIF1A C-terminal the DIDDI sequence that was previously shown to be important (35,40,48-50). In the archaeal case, aIF1A has neither DIDDI sequence nor a long C-terminal tail. However, an interaction involving the aIF5B archaeal-specific α15 helix and the C-terminal helix of aIF1A is observed. Moreover, a contact involving the 2-3 loop of aIF1A and the C-terminal helix of aIF5B is also visible, as in eukaryotes.

In the bacterial case, interactions between the IF1 and IF2 orthologs of e/aIF1A and e/aIF5B were also observed. However, these interactions strongly differ from those in the eukaryotic and archaeal cases. Indeed, domain IV of IF2 is shorter and does not contain the C-terminal helices (supplementary Figure 9). However, contact with IF1 is ensured by a bacterial-specific insertion in helix 8 of domain II (85-89,90, 91). Thus, interaction between e/aIF5B/IF2 and e/aIF1A/IF1 is universal but present domain-specific features that could have evolved lately. In all domains of life, the network of interactions involving the two initiation factors and the SSU would help to constrain the initiator tRNA in a conformation adequate for LSU joining, followed by the accommodation of the acceptor end in the PTC.

During eukaryotic subunit joining, it was proposed that rotation movements of the SSU with respect to the LSU controlled departure of eIF1A and GTP hydrolysis on eIF5B (34,45,46). The importance of ribosome dynamics during late steps of translation initiation was also illustrated in bacteria (90,60,76,84,86,92). The tRNA conformation observed in *P. abyssi* 30S:aIF1A:aIF5B:Met-tRNA_i_^Met^ is very close to that observed in *S. cerevisiae* 80S:eIF5B and reminiscent of the P/Pre-like state described in (45) (Figure 6C). Therefore, it is tempting to imagine that similar dynamics are also involved in the archaeal case. Nevertheless, further studies are clearly needed to understand how, in archaea, the 30S:aIF1A:aIF5B:Met-tRNA_i_^Met^ complex recruits the 50S in the presence of aIF1A and at which step GTP hydrolysis occurs. aIF5B release would leave a full ribosome in a semi-rotated state adequate for the recruitment of aEF1A:aa-tRNA and elongation. The present study paves the way for time-resolved cryo-EM using archaeal systems that should help answering these questions.

Finally, apart from its general role in canonical translation, eukaryotic eIF5B emerges as an important regulator of other specific nodes of translation such as ribosome biogenesis (93) or non-canonical translation initiation (94-96). Whether archaeal aIF5B is involved in similar processes remains an open and appealing question.

## Data availability

Cryo-EM maps have been deposited in the Electron Microscopy Data Bank under the accession numbers EMD-14731 (Map A, 30S:mRNA:ASL-Met-tRNA_i_^Met^), EMD-14579 (30S:mRNA:aIF1A:aIF5B-DIV:Met-tRNA_i_^Met^), EMD-14580 (Map B, IC3), EMD-14763 (Map B2, aIF1A:aIF5B:Met-tRNA_i_^Met^) and EMD-14581 (30S:mRNA:aIF1A:Met-tRNA_i_^Met^). The atomic models have been deposited to the Protein Data Bank under the accession numbers 7ZHG (30S:mRNA:ASL-Met-tRNA_i_^Met^), 7ZKI (aIF1A:aIF5B:Met-tRNA_i_^Met^), 7ZAH (IC3), 7ZAG (30S:mRNA:aIF1A:aIF5B-DIV:Met-tRNA_i_^Met^), 7ZAI (30S:mRNA:aIF1A:Met-tRNA_i_^Met^). X-ray data and models have been deposited with PDB IDs 7YYP (aIF5B:GDP) and 7YZN (aIF5B-ΔC:GTP).

## Acknowledgements

This work was supported by grants from the Centre National de la Recherche Scientifique and Ecole polytechnique to Unité Mixte de Recherche n°7654 and by a grant from the Agence Nationale de la Recherche (ANR-17-CE11–0037). Cryo-EM data benefited from access to the following facilities: eBIC, Diamond Light Source, UK; ESRF, Grenoble; Pasteur Institute, Paris and the Interdisciplinary Center for Electron Microscopy of Ecole Polytechnique (CIMEX). X-ray data were collected at Soleil synchrotron Proxima-1 and Proxima-2 beamlines. We thank the staff at these facilities, Alistair Siebert, Daniel Clare, Yuriy Chaban, David Owen (Diamond Light Source), Daouda Traoré, Michael Hons, Alessandro Grinzato (ESRF, Grenoble), Ileana Florea (CIMEX), William Shepard, Martin Savko (PX2) and Pierre Legrand (PX1) for their help with cryo-EM and X-ray data collections. We are particularly indebted to Stéphane Tachon (Institut Pasteur) for his help during the final cryo-EM data collection. We also thank Célia Plisson-Chastang (CBI, Toulouse) and Magali Blaud (CiTCom, Paris) for their kind gifts of cryo-EM grids at critical stages of this work and Sophie Bourcier (LCM, Ecole Polytechnique) for mass-spectrometry analyses.

