## Supplementary information for "Role of aIF5B in archaeal translation initiation"

| **Data collection** | aIF5B:GDP 7YYP | aIF5B-ΔC:GTP 7YZN |
|---|---|---|
| Molecule in a. u | 1 | 1 |
| Crystallization condition | 20 % PEG3350 ; 0.2M Lithium nitrate | 20% PEG3350 ; 8% Tacsimate pH=5 |
| Space group | $P2_1$ | $P4_12_12$ |
| Cell dimensions | | |
| a, b, c (Å) | 70.42 70.72 74.26 | 72.23 72.23 205.45 |
| α, β, γ (°) | 90 110.06 90 | 90 90 90 |
| Resolution (Å) | 48.31 - 2.9 | 49.59 - 1.7 |
| $R_{meas}$ | 0.169 (1.41)[a] | 0.111 (1.920) |
| I/σ(I) | 9.09 (1.12) | 20.16 (1.88) |
| Completeness (%) | 99.66 (98.42) | 99.5 (97.0) |
| Redundancy | 6.9 (6.0) | 26.3 (25.0) |
| $CC_{1/2}$[b] | 0.997 (0.672) | 0.999 (0.712) |
| Unique reflections | 15340 (1498) | 61785 (9577) |
| **Refinement** | | |
| $R_{work}/R_{free}$[c] | 0.213/0.263 | 0.169/0.197 |
| No atoms | 4783 | 4006 |
| Protein | 4700 | 3627 |
| Waters | 35 | 345 |
| Heterogen atoms | 48 | 34 |
| B-factors (Å$^2$) protein | 78.2 | 40.5 |
| Nucleotide/$Mg^{2+}$/Na | 58.2 | 19.9/19.1/31.7 |
| Waters | 35 | 42 |
| Bond lengths (Å) | 0.02 | 0.011 |
| Bond angles (°) | 0.54 | 1.12 |

**Supplementary Table 1: Crystallographic structures, data collection and refinement statistics**
A single crystal was used for data collection.
[a] Values in parentheses are for highest-resolution shell.
[b] $CC_{1/2}$ is the correlation coefficient between two random half data sets (1).
[c] $R_{free}$ is calculated with 5% of the reflections.

| | 30S-high-res *Map A* | aIF1A:aIF5B:Met-tRNA$_i^{Met}$ *Multibody map B2* | IC3 *Map B* | 30S:mRNA:aIF1A:aIF5B-DIV:Met-tRNA$_i^{Met}$ | 30S:mRNA:aIF1A:Met-tRNA$_i^{Met}$ |
|---|---|---|---|---|---|
| **Data collection and processing** | | | | | |
| EMD number | EMD-14731 | EMD-14763 | EMD-14580 | EMD-14579 | EMD-14581 |
| PDB number | 7ZHG | 7ZKI | 7ZAH | 7ZAG | 7ZAI |
| Microscope | TFS Krios | | | | |
| Camera | Gatan K3 with Bioquantum | | | | |
| Magnification | 105 000x | | | | |
| Voltage (kV) | 300 | | | | |
| Electron exposure (e–/Å$^2$) | 40 | | | | |
| Defocus range (µm) | -0.8 – -3 | | | | |
| Pixel size (Å) | 0.86 | | | | |
| Symmetry imposed | C1 | | | | |
| Initial particle images (no.) | ~2,000,000 | | | | |
| Final particle images (no.) | ~1,100,000 | ~37,000 | ~37,000 | ~193,000 | ~382,000 |
| Resolution (unmasked, Å) | 2.5 | - | 3.2 | 3.0 | 2.9 |
| Resolution (masked, Å) | 2.25 | 3.6 | 2.7 | 2.8 | 2.6 |
| FSC threshold | 0.143 | 0.143 | 0.143 | 0.143 | 0.143 |
| **Refinement** | | | | | |
| Initial model used (PDB code) | 6SWC | 7YYP, 7YZN | 7ZHG, 7YYP, 7YZN | 7ZHG,7YYP, 7YZN | 7ZHG |
| d FSC model (Å), threshold 0.5 | 2.3 | 4.2 | 2.7 | 2.9 | 2.7 |
| Map sharpening *B* factor (Å$^2$) | -74 | -82 | - | - | - |
| **Model composition** | | | | | |
| Non-hydrogen atoms | 64,278 | 7,120 | 69,356 | 65,678 | 64,649 |
| Protein residues | 3,601 | 687 | 4,287 | 3,823 | 3,693 |
| Nucleotides | 1,502 | 76 | 1,564 | 1,564 | 1,564 |
| Ligands | 90xMg, 6xZn | 1xMg, 1xGNP | 63xMg, 6xZn | 62xMg, 6xZn | 62xMg, 6xZn |
| Water molecules | 2083 | 0 | 379 | 379 | 379 |
| **Average *B* factors (Å$^2$)** | | | | | |
| Protein | 16.9 | 52.2 | 68.6 | 96.6 | 89.1 |
| Nucleic acid | 19.3 | 108.9 | 59.2 | 113.5 | 91.1 |
| Ligand | 13.5 | 72.8 | 45.5 | 114.6 | 85.2 |
| Water molecules | 11.1 | - | 37.9 | 82.2 | 66.5 |
| **R.m.s. deviations** | | | | | |
| Bond lengths (Å) | 0.004 | 0.002 | 0.008 | 0.006 | 0.004 |
| Bond angles (°) | 0.755 | 0.552 | 0.707 | 0.567 | 0.555 |
| **Validation** | | | | | |
| MolProbity score | 1.40 | 1.54 | 1.45 | 1.39 | 1.29 |
| Clashscore | 2.83 | 9.39 | 3.72 | 3.94 | 3.40 |
| Poor rotamers (%) | 1.41 | 0.17 | 1.24 | 0.71 | 0.64 |
| **Ramachandran plot** | | | | | |
| Favored (%) | 96.67 | 97.8 | 96.59 | 96.70 | 97.08 |
| Allowed (%) | 3.33 | 2.20 | 3.41 | 3.30 | 2.92 |
| Disallowed (%) | 0.00 | 0.00 | 0.00 | 0.00 | 0.00 |
| **Correlation coefficients** | | | | | |
| Mask CC | 0.86 | 0.73 | 0.92 | 0.91 | 0.94 |
| Volume CC | 0.83 | 0.70 | 0.91 | 0.90 | 0.93 |

**Supplementary Table 2: Cryo-EM data collection, refinement and validation statistics**

| **Modification Name** (PDB identifier) | | **Modification type** | **Position(s) in 16S rRNA sequence of *P. abyssi*** |
|---|---|---|---|
| **ac4C** | (4AC) | N4-acetylcytidine | 17, 53, 286, 303, 319, 379, 394, 479, 511, 546, 590, 626, 636, 703, 718, 731, 751, 828, 839, 848, 851, 868, 957, 1028, 1147, 1233, 1239, 1479 |
| **ac4Cm** | (LHH) | N4-acetyl-2'-O-methylcytidine | 250, 1041 |
| **Am** | (A2M) | 2'-O-methyladenosine | 373 |
| **Cm** | (OMC) | 2'-O-methylcytidine | 129, 846, 1036, 1040, 1376 |
| **Gm** | (OMG) | 2'-O-methylguanosine | 467, 471, 519, 657, 680, 873, 913, 934, 1069 |
| **m1Y** | (B8H) | 1-methylpseudouridine | 938 |
| **m3U** | (UR3) | 3-methyluridine | 1467 |
| **m5C** | (5MC) | 5-methylcytidine | 535, 693, 875, 1025, 1202, 1374, 1496, 1498, 1505 |
| **m6A** | (6MZ) | N6-methyladenosine | 1469 |
| **m6,6A** | (MA6) | N6,N6-dimethyladenosine | 1487, 1488 |
| **Um** | (OMU) | 2'-O-methyluridine | 20, 64, 774, 787, 830, 1177, 1380 |
| | | Unidentified modification | 939,1378 |

**Supplementary Table 3** : **Modified residues identified in 16S rRNA sequence of *P. abyssi*.**
The short name, the PDB identifier, the name and the position in 16S rRNA sequence of *P. abyssi* are indicated for each type of modification. In IC2 (PDB ID 6SWC), residue C939 had been modeled as an m5C on the basis of additional density and of the presence of an ortholog of the corresponding modification enzyme (RsmB) in the *P. abyssi* genome. However, the 2.25 Å resolution map A does not confirm this modification but rather shows an extra density on N4. Because the corresponding modification could not be identified unambiguously, residue 939 was modeled as C. Similarly, extra density for C1378 is clearly visible, however the corresponding modification could not be identified. Four ac4C (648, 998, 1184, 1193) previously modeled in IC2 and confirmed by reverse transcription (2) were not modeled in IC3 because of insufficient density. This discrepancy may be linked to the dynamic character of ac4C modifications (3).

| aIF5B | rRNA | Type | Conservation Archaea | Conservation Eukarya |
|---|---|---|---|---|
| R280 (sc) DII | U377 (phosphoryl) | H-bond | 65% R; ~100 % K or R | 99.2% R; 100% K or R |
| R280 (sc) DII | U377 (ribose and base) | Hydrophobic | See above | See above |
| Y300 (sc) DII | OMU64 | OMe-aromatic | Not conserved | Not conserved |
| P245-L247 (TTT tight turn) DII | A63 (ribose and base) h5 | Hydrophobic | 95% G246 ; 69% L247 (85% L or F) | 100% G246 ; 53% L247 (84% L or F) |
| G246-L247 DII | G367 (ribose) h5 | Hydrophobic | See above | See above |
| K392 (sc) DIII | G371 (phosphoryl) h14 | H-bond | 47% K ; 81% K or R | 93% K; ~100% K or R |
| D421 (sc) DIII | C370 (phosphoryl) | H-bond | 56% D ; 78% D or E | 27% D ; ~100% D or E |
| **aIF5B** | **uS12** | | | |
| R383 (sc) | H100 (mc) | H-bond | 50% K ; 65% K or R | Very rarely K or R |
| E404 (sc) | H100 (sc) | H-bond | Not conserved (15%, often deletion) | Majorily K, R or H |
| K392 (sc) | I78 (mc) | H-bond | See above | See above |
| M396 (sc) | I53 (sc) | Hydrophobic | 12%M; 79% M,L,V,I,A | 79% M; 98% M,L,V,I |
| E397 (sc) | I53 (sc) | Hydrophobic | 47% E ; 60% D or E | Never E or D; 69% K or R. |
| T393 (sc) | I78 (sc) | Hydrophobic | Not conserved (5%), rather R or K (81%) | not conserved, rather K or R (97%). |
| L399 (sc) | Q76 (sc) | Hydrophobic | Not conserved (5%), rather S (28%) | Almost never L, rather S (42%) |
| S400 (sc) | L55 (sc) | Hydrophobic | 14% S; 42% S or T | 1% S; 48% S or T |
| V401 (sc) | H100 (sc) | Hydrophobic | 2% V, 43% V,L,M,I,A | 0.1% V, rather M (97.4%) |
| **aIF5B** | **tRNA** | | | |
| F481 (sc) | esterified Met (mc) | Hydrophobic | Close to 100% F | 89% F; 100% F,I,Y,V |
| Y479 (sc) | esterified Met | Hydrophobic | 33% Y ; 71% Y,F,H | 2% Y ; 7% Y,F,H |
| 517-520 ($\beta$27) | Met-A76 | Hydrophobic | Pos. 518: 66% I; ~100% I,V,M,L | Pos. 518 : 69% I; ~100% I,V,M,L |
| 533-537 ($\beta$30) | Met-A76 | Hydrophobic | 534-537 conserved in chemical character with a VAVS consensus | VAV consensus at 534-536, but at 537 87% K (close to 100% R or K) |
| I488 (sc) | A76 (base) | Hydrophobic | Close to 100% I or V, rarely A or L | 99% I or V, rarely T or L |
| F481 (sc) | A76 (ribose) | Pi-interaction | See above | See above |
| F481 (mc) | C75 (mc) | Hydrophobic | See above | See above |
| R482 (sc) | C75 (mc) | H-bond | 99% R | Mostly N (81%). |
| F481 (mc) | C74 (ribose) | H-bond | See above | See above |
| R482 (sc) | C74 (base) | Pi-interaction | See above | See above |
| R482 (sc) | C74 (ribose) | Hydrophobic | See above | See above |
| R483 (mc) | C74 (base) | Hydrophobic | 24% R; 3% K; 40% Q; | 2% R; 87%K; 1% Q |
| S484 (sc) | C74 (base) | H-bond | Mostly S (65%) or N (20%) | Mostly K or R (80%) |
| S484 (mc) | C74 (base) | Hydrophobic | See above | See above |
| R483 (mc) | A73 (base) | Hydrophobic | See above | See above |
| R545 (sc) | G70 (base) | H-bond | 92% R. K in a few taxons. | 99% R ; K in a few taxons. |
| R545 (sc) | G70 (phosphoryl) | H-bond | See above | See above |
| R483 (sc) | A1 (base) | Pi-interaction | See above | See above |
| R545 (sc) | C69 (base) | PI-interaction | See above | See above |

**Supplementary Table 4: Main interactions of aIF5B in the IC3 complex.**
Interactions were computed using LigPlot+ (4) and verified manually by checking the cryo-EM maps B and B2. Conservation of the concerned aIF5B residue in archaeal and eukaryal sequences is indicated. We used an alignment of 2957 archaeal and 2143 eukaryal sequences performed with ClustalX (5) and refined it manually. Note that the same alignment showed that $\alpha$15 of aIF5B, although not conserved in sequence, is almost systematically present in archaeal sequences.

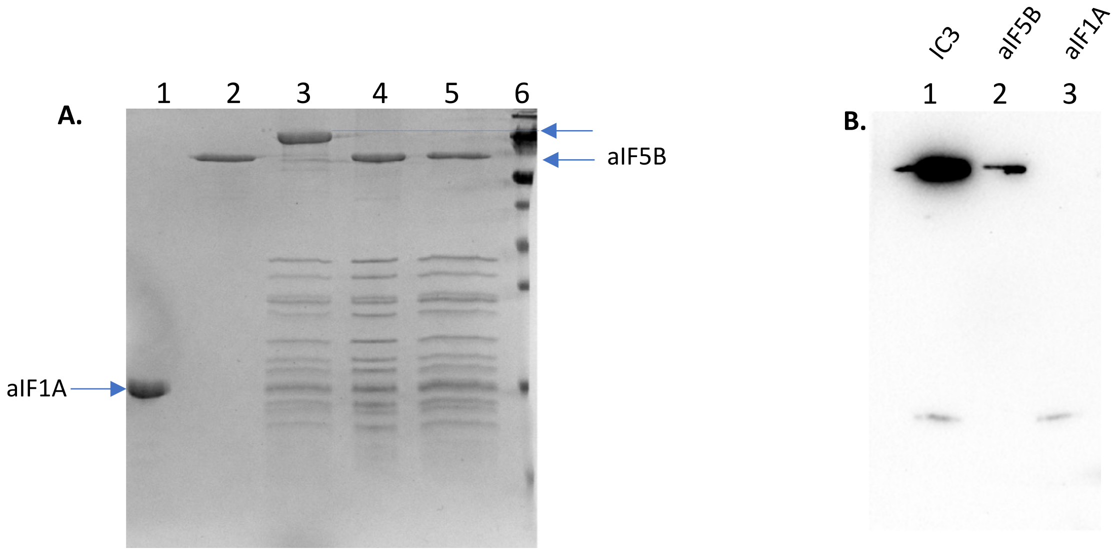

**Supplementary Figure 1: Purification of IC3 from *P. abyssi* analyzed by SDS-PAGE and Western blot**
(**A**) SDS-PAGE analysis of IC3 purification steps.
Lane 1: purified N-terminally tagged version of aIF1A.
Lane 2: purified N-terminally tagged version of aIF5B.
Lane 3: purified 30S subunits from *P. abyssi*. The band corresponding to contaminating phosphoenol pyruvate synthase (Pep synthase) is indicated by an arrow. This band is removed upon affinity purification.
Lane 4: IC3 after affinity column purification and concentration. The band corresponding to aIF1A co-migrate with ribosomal proteins. The presence of tRNA was verified using staining with ethidium bromide (not shown).
Lane 5: IC3 after buffer exchange before $BS^3$ crosslinking.
Lane 6: molecular weight marker (LMW, 70, 55, 45, 35, 25, 15, 10 kDa).
(**B**) Detection of aIF5B and aIF1A in IC3 by western blotting using antibodies directed against their His-tag.

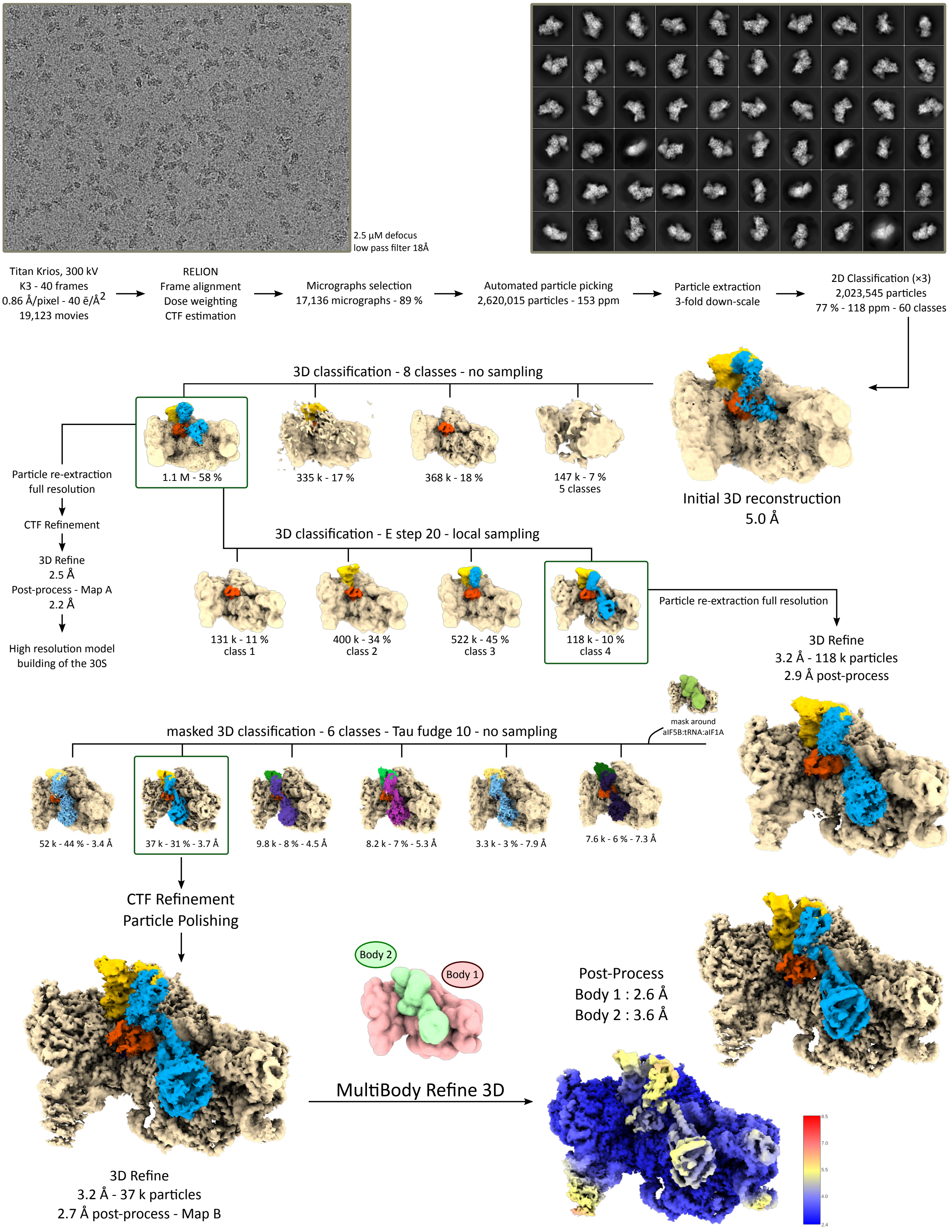

**Supplementary Figure 2: Cryo-EM data processing flowchart.**
Data processing was performed using RELION (6,7). The initiation factors bound to the 30S subunit (beige) are colored as follows: aIF5B blue, aIF1A orange and tRNA yellow. Several steps of 3D classification allowed us to identify a subset of 118 k particles yielding clear density for all the elements of the complex. Further 3D focused classification sorted out remaining heterogeneity and identified a final pool of 37k particles. CTF Refinement and particle polishing improved the resolution to 2.7 Å. The multibody refinement maps shown are composite maps from the two independently refined bodies with their respective estimated resolution after post-processing. They are colored with the usual color scheme (top) and by local resolution (bottom) (8).

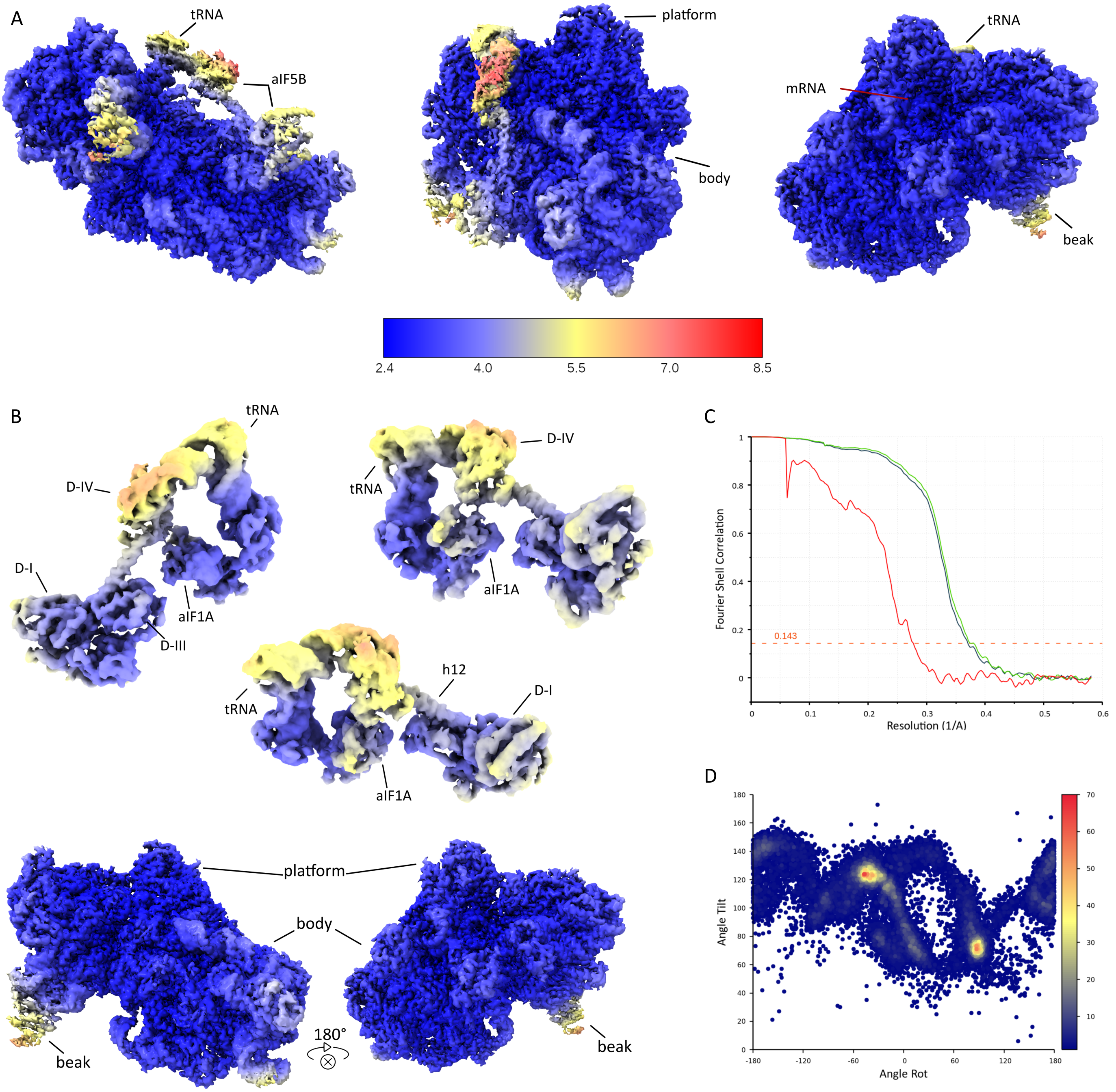

**Supplementary Figure 3: Local resolution, Fourier shell correlation (FSC) and particle distribution of the final maps.** (**A**) Three views of the final full map colored by local resolution values (key shown). The head and body of the 30S subunit are at the same resolution confirming that there is no head movement. Local resolution of domain II and domain III of aIF5B is close to that of the 30S subunit whereas peripheral regions of the factor are defined at lower resolution, pointing to some residual heterogeneity. Particle distribution is shown as a colored 2D-projection of distribution histograms as a function of Rot and Tilt angles.

(**B**) Local resolutions of the two multi-body maps showed from multiple viewing angles. The same color scale is used for all maps. After multi-body refinement the local resolution for domain-IV and the acceptor helix of the Met-$tRNA_i^{Met}$ is greatly improved.

(**C**) FSC curves of the full map (map A, blue), and the two multibody maps (green and red, maps B1 and B2) are drawn on the same graph. The 0.143 Gold-standard threshold is shown as an orange doted line.

(**D**) Particle distribution is shown as a colored 2D-projection of distribution histograms as a function of Rot and Tilt angles.

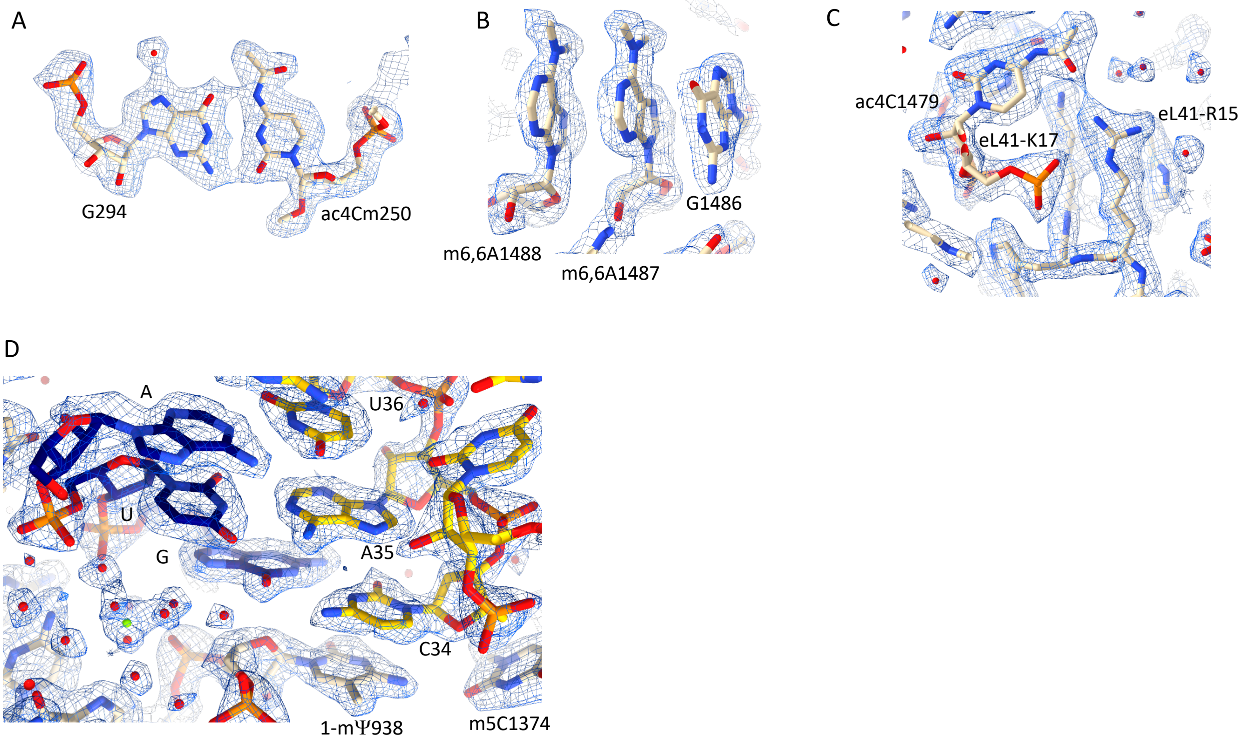

**Supplementary Figure 4: Cryo-EM map A in some regions of the 30S.**
(**A**) Example of an N4-acetyl-2'-O-methylcytidine.
(**B**) The two di-methyl adenosines 1488 and 1487 are shown.
(**C**) Interaction of ac4C1479 with eL41-R15.
(**D**) Codon:anticodon interaction and surrounding residues. Water molecules are shown as red spheres and the hexacoordinated magnesium ion is green.

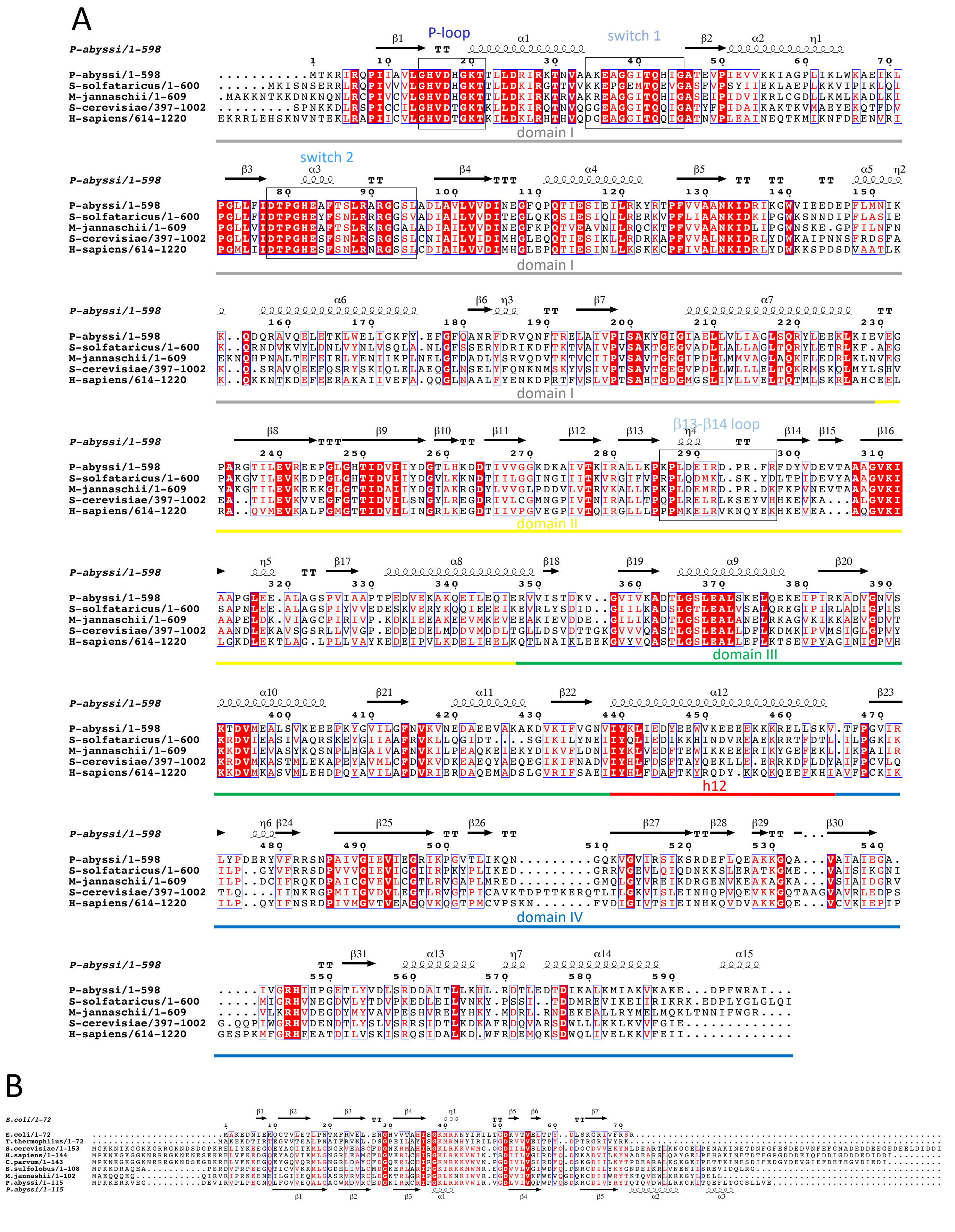

**Supplementary Figure 5: e/aIF5B and e/aIF1A.**

(**A**) Multiple sequence alignment of archaeal and eukaryotic e/aIF5B manually refined according to structural data. The figure was drawn with Espript (9).

(**B**) Multiple sequence alignment of archaeal and eukaryotic e/aIF1A refined according to structural data. The sequence of the *E. coli* IF1 orthologue is also shown. The alignment is from (10).

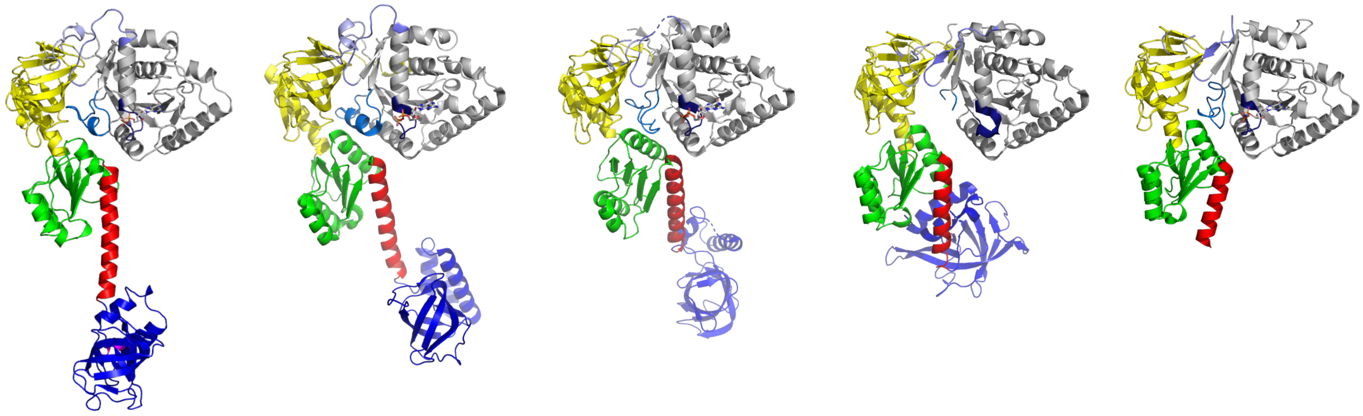

| | 5FG3 (GDP) | 1G7S (GDP) | 3WBI (APO) | 4NCL (GDP) |
|---|---|---|---|---|
| rmsd between domains I | 0.6 Å (178 atoms) | 0.72 Å (166 atoms) | 0.98 Å (166 atoms) | 0.89 Å (163 atoms) |
| Movement of DII | 9.2°, 1.7 Å | 11.6°, 2 Å | 5.6°, 1 Å | 9°, 2.3 Å |
| Movement of DIII | 22°, 4 Å | 35°, 7.4 Å | 15°, 6.5 Å | 17.6 °, 4.2 Å |

**Supplementary Figure 6: Structural alignments of e/aIF5B in switch OFF states.**
The structure of aIF5B from *P. abyssi* was chosen as a reference. The structures (PDB IDs 5FG3 (11), 1G7S (12), 3WBI (13), 4NCL (14)) were superimposed on domain I of aIF5B. The color code is the same as in Figure 2 .
Rmsd were calculated with Pymol. Movements of DII and DIII (rotation, translation) were calculated with Pymol using the script https://raw.githubusercontent.com/speleo3/pymol-psico/master/psico/orientation.py.

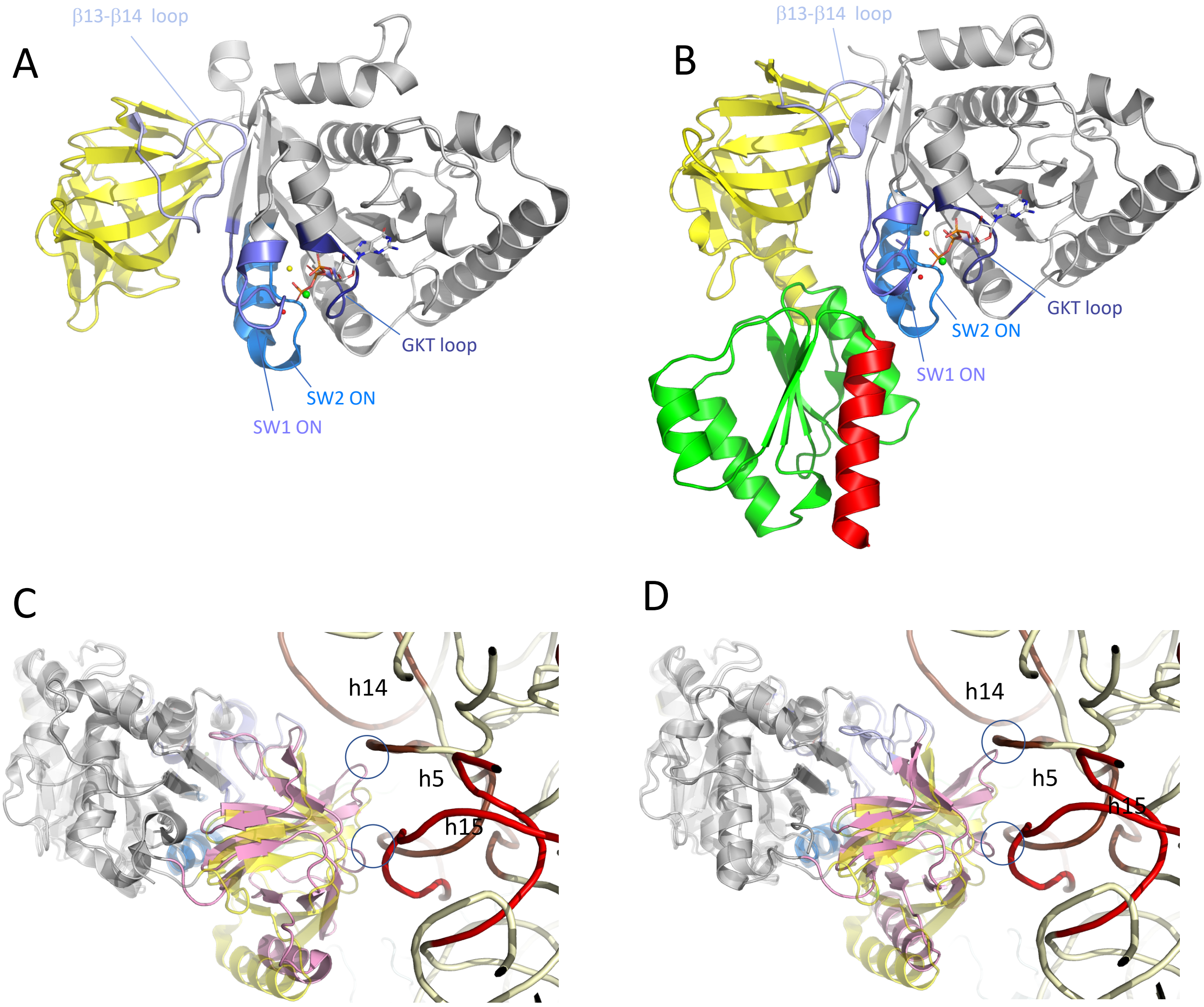

**Supplementary Figure 7: Comparison of Pa-aIF5B with Ct-aIF5B.**

(**A**) Cartoon representation of Ct-DI-DII-eIF5B:GTP (PDB ID 4TMX (14)). The color code is the same as in Figure 1. Domain I of Ct-DI-DII-eIF5B:GTP was superimposed onto domain I of aIF5B-ΔC:GTP with an rmsd of 0.575 Å for 190 atoms compared.

(**B**) Cartoon representation of aIF5B-ΔC:GTP. Panels A and B show that the relative orientations of domains I and II are close in the two structures.

(**C**) Domains I of Ct-DI-DII-eIF5B:GTP and aIF5B in IC3 were superimposed. The color code is the same as in Figure 2 except that Ct-DI-DII-eIF5B:GTP domain II is in light pink. Ct-DI-DII-eIF5B:GTP is shown at the foreground and aIF5B in IC3 is shown using transparent cartoons as a reference.

(**D**) Domains I of aIF5B-ΔC:GTP and aIF5B in IC3 were superimposed. aIF5B-ΔC:GTP domain II is in pink. aIF5B-ΔC:GTP is shown at the foreground and aIF5B in IC3 is shown using transparent cartoons as a reference (same as Figure 5D shown here for comparison). The C and D views show that the position of domain II observed in aIF5B-ΔC:GTP or in Ct-DI-DII-eIF5B:GTP would create clashes with h5 and h15 (blue circles).

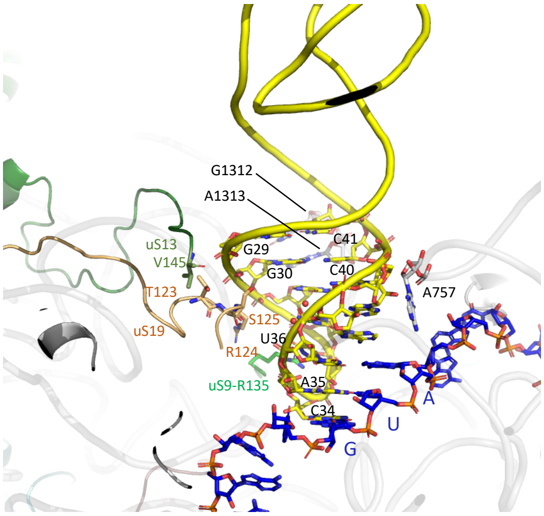

**Supplementary Figure 8: Initiator tRNA interactions at the P site.**
uS13 (dark green), uS19 (light orange) and uS9 (green) C-terminal tails and residues described in text are shown. Bases G1312 and A1313 interacting with the G29-C41 and the G30-C40 base pairs of the Met-tRNA$_i^{Met}$ anticodon stem as well as A757 are shown in sticks.

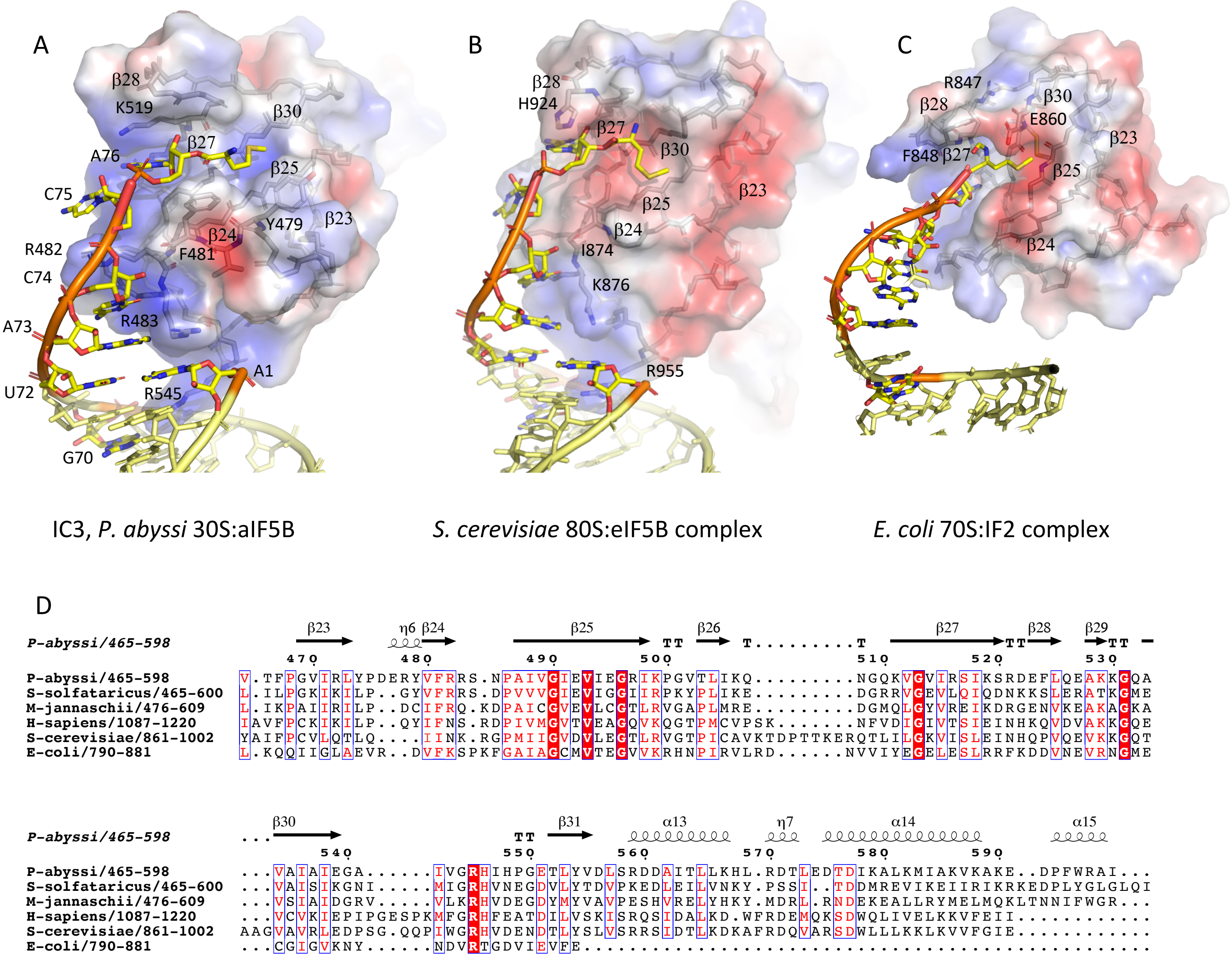

**Supplementary Figure 9: Binding of the CCA-end of the initiator tRNA**

The electrostatic potential surfaces of e/aIF5B-IF2-DIV bound to initiator tRNA are shown for *P. abyssi* (**A**), *S. cerevisiae* (PDB ID 6WOO (15)) (**B**) and *E. coli* (PDB ID 3JCJ (16)) (**C**) Main residues involved in interaction with the initiator tRNA are shown. In panel C, R847, F848, E860 are involved in the recognition of the formyl group. Residues C815 and C861 that stabilize the binding pocket through a disulfide bond are also shown. These residues are widely conserved in bacteria. (**D**) Multiple sequence alignment of domains IV of a/eIF5B and *E. coli* IF2.

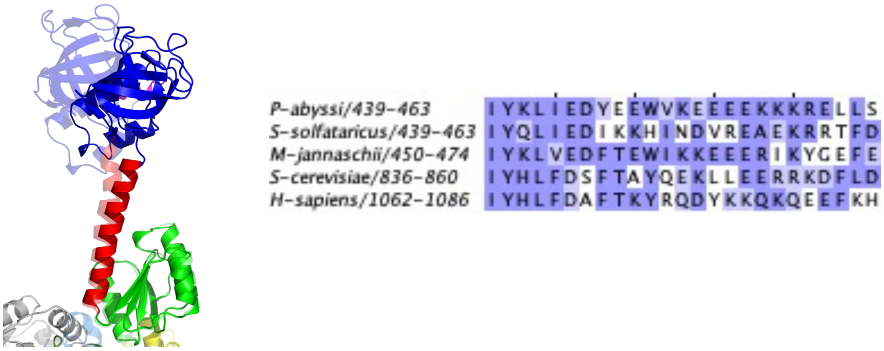

**Supplementary Figure 10: Orientation of h12 in aIF5B:GDP and in IC3.**

Domains III of the two structures were superimposed. aIF5B:GDP is shown at the foreground and aIF5B in IC3 is shown using transparent cartoons as a reference. The view shows the bending of the h12 helix originating in the middle of it, at a sequence very rich in hydrophilic residues. Stretches of lysine and glutamate residues are found in eukaryotes and archaea. DIV can rotate freely with respect to helix 12, a motion originating from the region of K464 (see also supplementary Figure 5).
